# Elevator mechanism dynamics in a sodium-coupled dicarboxylate transporter

**DOI:** 10.1101/2022.05.01.490196

**Authors:** Colin D. Kinz-Thompson, Maria Louisa Lopez-Redondo, Christopher Mulligan, David B. Sauer, Jennifer J. Marden, Jinmei Song, Emad Tajkhorshid, John F. Hunt, David L. Stokes, Joseph A. Mindell, Da-Neng Wang, Ruben L. Gonzalez

**Affiliations:** Department of Chemistry, Columbia University, New York, New York 10027, US; Department of Cell Biology, New York University School of Medicine, New York, New York 10016, US; School of Biosciences, University of Kent, Canterbury, Kent, CT2 7NH, UK; Theoretical and Computational Biophysics Group, NIH Center for Macromolecular Modeling and Bioinformatics, Beckman Institute for Advanced Science and Technology, Department of Biochemistry, Center for Biophysics and Quantitative Biology, University of Illinois at Urbana-Champaign, Urbana, IL 61801, US; Department of Biological Sciences, Columbia University, New York, New York 10027, US; Membrane Transport Biophysics Section, Porter Neuroscience Research Center, National Institute of Neurological Disorders and Stroke, National Institutes of Health, Bethesda, MD 20892, US

## Abstract

VcINDY, the sodium-dependent dicarboxylate transporter from *Vibrio cholerae*, is responsible for C_4_- and C_5_-carboxylate uptake into cells. The molecular mechanism of how VcINDY physically moves substrates across the membrane, and does so in an energetically efficient manner, is unclear. Here, we use single-molecule fluorescence resonance energy transfer experiments to directly observe the individual mechanistic steps that VcINDY takes to translocate substrates across a lipid bilayer, and then test key predictions of transport cycle mechanistic models. Our data provide the first direct evidence that VcINDY undergoes stochastic, elevator-type conformational motions that enable substrate translocation. Kinetic analysis suggests that the two protomers of the VcINDY homodimer undergo those motions in a non-cooperative manner, and thus catalyze two independent transport reactions. The relative substrate independence of those motions supports the notion that the VcINDY transport cycle maintains strict co-substrate coupling using a cooperative binding mechanism. Finally, thermodynamic modeling provides insight into how such a cooperative binding mechanism provides a generalized approach to optimizing transport for many secondary active transporters.

**Significance Statement:** Transporter proteins use energy to move molecular materials into and out of cells. To be efficient, the transporter motions responsible for moving the molecules must be tightly choreographed to avoid wasting energy without transporting anything. By measuring the motions and kinetics of a prototypical transporter (VcINDY) at the single-molecule level, this study finds the first evidence that transporters like VcINDY achieve efficient transport by coordinating constantly dynamic, “elevator-type” motions while sitting in the cellular membrane. The efficiency of these surprisingly dynamic transporters is then revealed by thermodynamic modeling, which explains the molecular basis behind how highly cooperative, substrate binding reactions may have evolved as the optimal strategy for maximizing transporter efficiency.

## Introduction

Secondary active transporters are transmembrane proteins that harvest free energy from electrochemical gradients to drive the movement of substrates across a membrane (Fig. 1) (1). In the absence of a free energy source, these transporters move substrates back and forth across the membrane without creating a net change in substrate concentration (2). Net transport occurs because the transporter can dissipate the electrochemical gradient of a ‘driving’ substrate to harvest free energy, and then couple that free energy to the ‘uphill’ movement of a ‘driven’ substrate across the membrane (3). The mechanistic details of this coupling are governed by the transport cycle, which describes how the transporter binds its substrates, translocates them across the membrane, releases them, and resets itself for the next round (Fig. 1B). While the specific details of the transport cycle and how it is achieved on a molecular level are unique to each transporter or transporter family, the active use of free energy suggests that transport cycles must maintain strict substrate coupling to minimize energetically wasteful ‘slippage cycles’ (*c*.*f*., Fig. 1B) in which driving substrates are transported without driving the translocation of the driven substrate (1).

**Figure 1.**
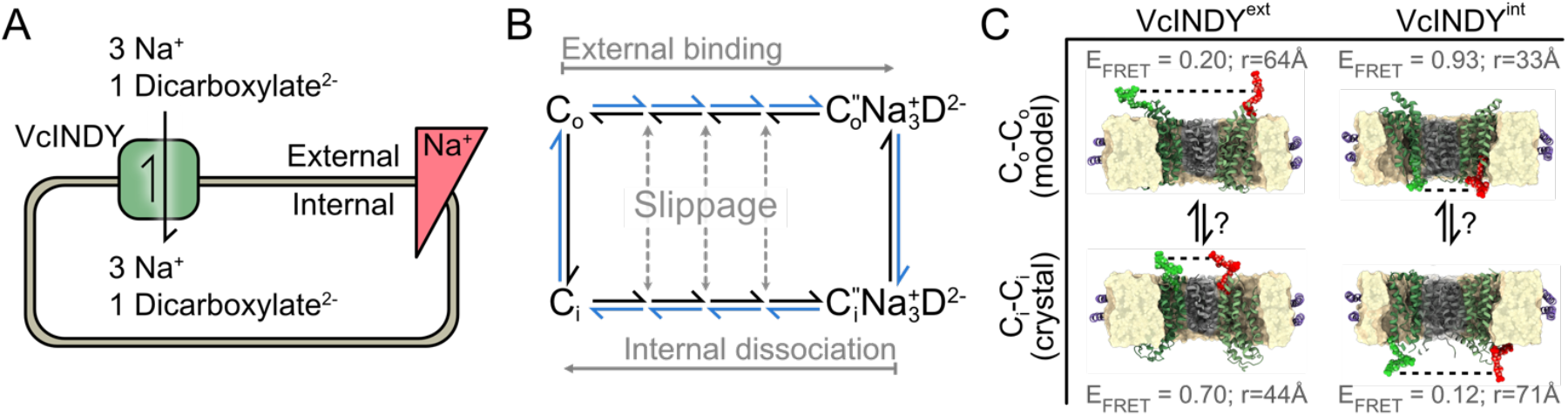
VcINDY transport cycle. (A) Diagram of VcINDY dimer (green) reversibly transporting sodium ions (Na^+^) and dicarboxylate substrate (D^2-^) into and out of a cell with a stoichiometry of 3:1; an inward facing Na^+^ electrochemical gradient provides the free energy for vectorial transport. (B) Minimal mechanistic model of VcINDY transport cycle. Outward- and inward-facing conformations of the VcINDY homodimer (C_o_, and C_i_, respectively) bind Na^+^ and D^2-^ in an unknown order along the horizontal transitions; double-prime signifies both co-substrates are bound. VcINDY protomer conformational motions along the vertical transitions are involved in substrate translocation. At least one closed transport cycle, such as shown here in the clockwise direction in blue, is required for vectorial transport. Slippage reactions occur when an incomplete set of co-substrates is moved across the membrane. (C) Cartoon diagram of smFRET pairs designed to report on the VcINDY translocation reaction for VcINDY^ext^ (left; G211) and VcINDY^int^ (right; S436). The distance between each labeling position, *r*, was determined from the C_α_-C_α_ distance, and was measured for a C_o_, substrate-bound repeat-swap model (C_o_-C_o_, PMDB: PM0080216), and a substrate-bound, C_i_ X-ray crystallography structure (C_i_-C_i_, PDB: 5UL9). E_FRET_ was estimated using R_0_=51 Å. Scaffold domains are shown in gray, and transport domains are shown in green.

While all secondary active transporters work by facilitating a general, dynamic equilibrium (*e*.*g*., *mA*_outside_+ *nB*_outside_⇌ *mA*_*inside*_ + *nB*_*inside*_, where *A* and *B* are co-substrates with stoichiometries *m* and *n*), the molecular details of how this chemical reaction maps onto the transport cycle are determined by the conformational dynamics of the transporter. According to the alternating access hypothesis, secondary active transporters must undergo a reversible conformational transition between at least two distinct conformational states: one with the substrate binding pocket in a state that is ‘outward-facing’ with respect to the membrane (C_o_), and one in an ‘inward-facing’ state (C_i_) (2). While there are many possible structural implementations of alternating access, including the so-called ‘rocker-switch’, ‘rocking-bundle’, and ‘elevator’ transport mechanisms (4), at least two distinct conformational transitions between C_i_ and C_o_ must occur in a complete transport cycle (*c*.*f*., Fig. 1B). Here, we investigate the role of such conformational dynamics in the transport cycle of the prototypical sodium-dependent divalent anion transporter from mesophilic *Vibrio cholerae*, VcINDY, and elucidate how they are tuned to enable efficient transport and minimize slippage.

VcINDY is a well-characterized member of the divalent anion/sodium symporter (DASS) class of secondary active transporters, which extends across all kingdoms of life. In mammals, it is represented by the solute carrier 13 (SLC13) gene family, which includes members that transport di- and trivalent anions, including sulfate, citrate, and succinate (5, 6). Within this family, the SLC13A5 citrate transporter, NaCT, has significant metabolic and neurologic health importance for humans, and mutations in NaCT can cause epilepsy in children (reviewed in (7)). Like NaCT, VcINDY is a Na^+^-driven carboxylate transporter, but VcINDY is primarily a succinate transporter with an electrogenic transport cycle stoichiometry of 3 Na^+^ ions per succinate (Fig. 1A) (8, 9). Over the past decade, extensive *in vitro* biochemical characterization (8, 10, 9, 11, 12), and deep structural insight from X-ray crystallography, cryo-electron microscopy (cryo-EM), and computational studies (13, 10, 14–16), have made VcINDY into the prototypical transporter for understanding the mechanistic details of transport in the DASS family. While the biochemical and structural studies published to-date do not yield a description of the molecular mechanism for the complete VcINDY transport cycle, recent combinations of computational modeling, biochemical, and structural analysis have provided insight (10, 17). Structural studies have revealed VcINDY is a homodimeric membrane protein with two-fold rotational symmetry; however, in all these studies, VcINDY was only observed in a single C_i_ conformational state in which the substrate binding pockets of both protomers are accessible only from the cytosolic side (*i*.*e*., the global homodimer conformation is C_i_-C_i_) (13, 14, 17, 18). Computational and crosslinking studies suggest that VcINDY functions using an elevator mechanism, with a large translocation of the ‘transport’ domain that carries substrates from one side of the membrane to the other, and a relatively stationary ‘scaffold’ domain that harbors the dimerization interface (10). This model is supported by recent structural work revealing a C_o_ state for LaINDY, a close homolog from *Lactobacillus acidophilus* (17). While these lines of evidence support an elevator mechanism, there has been no experimental observation of a transition between these C_o_ and C_i_ states, nor any direct evidence that they are actually responsible for achieving alternating access during transport. Comprehensive validation of this model requires confirming the proposed elevator motions, connecting them to the transport cycle, and understanding how those conformational dynamics enable VcINDY to achieve tight co-substrate coupling between Na^+^ and succinate to minimize energetically costly slippage of substrates across the membrane (Fig. 1B).

In this work, using single-molecule fluorescence resonance energy transfer (smFRET) microscopy, we aimed to directly observe the global conformational dynamics of homodimeric VcINDY in a lipid-bilayer membrane, allowing us to characterize the structural and kinetic details of its transport cycle. To achieve this, we designed two complementary VcINDY constructs for smFRET experiments. Each smFRET construct contained a FRET donor- and acceptor fluorophore pair engineered at residue positions such that the resulting donor-acceptor distance-dependent FRET efficiency (E_FRET_) would report on the conformational dynamics responsible for substrate translocation (Fig. 1C). These smFRET constructs can therefore be used to directly observe the motions that VcINDY undergoes to achieve alternating access (Fig. 1B). In performing smFRET experiments with these constructs, we obtained direct experimental evidence that (*i*) global conformational dynamics of VcINDY during the transport cycle are consistent with the elevator model of transport, (*ii*) the individual protomers of homodimeric VcINDY exhibit minimal cooperativity, and (*iii*) VcINDY minimizes slippage using a mechanism in which co-substrate binding is highly cooperative. These findings provide the first comprehensive description of the molecular underpinnings of the VcINDY transport cycle, and, by extension, deep insight into the transport cycle of all DASS transporters.

## Results

### smFRET constructs reporting on substrate translocation were designed, constructed, and functionally validated

In order to directly monitor the VcINDY conformational dynamics relevant to the transport cycle (Fig. 1B), we designed two smFRET constructs based on the assumption that substrate translocation in VcINDY involves a large-scale conformational change between the C_o_-C_o_and the C_i_-C_i_ conformations (*c*.*f*., Fig. 1C) (10, 13, 14, 17). One construct was designed to contain a FRET donor- and acceptor fluorophore pair on the external, periplasmic face of the transport domains of VcINDY whereas the other construct was designed to contain them on the internal, cytosolic face. These complementary designs comprise a ‘multi-perspective’ approach with an internal self-consistency where any global conformational motions of the VcINDY homodimer occurring during the transport cycle (*e*.*g*., an elevator motion) can be detected on both the external and internal sides of the protein.

To design these ‘external’ and ‘internal’ smFRET constructs, we identified solvent-exposed residues in the transport domain with optimal distances between the two equivalent positions across the homodimer (Fig. 1C). Specifically, residues were chosen such that the distance would undergo a significant change upon a transition between the structurally defined C_i_-C_i_ and the computationally predicted C_o_-C_o_states (10, 13, 14, 17). In an optimized smFRET experiment, the corresponding change in the distance-dependent E_FRET_ will utilize the full dynamic range of E_FRET_ from zero to one. To achieve this, we sought residues that moved from separations of less than the half-maximum E_FRET_ of the donor-acceptor pair (R_0_) to more than the R_0_, or vice versa. We used Alexa Fluor 555 as our donor and Alexa Fluor 647 as our acceptor, with an R_0_ ∼51 Å (19). Given these considerations, we chose Gly211, located on the extracellular face of the transport domain as our external site, with a predicted E_FRET_ change from ∼0.20 to ∼0.93 upon a transition from the C_o_-C_o_to the C_i-_C_i_ state (Fig. 1C). Likewise, we chose Ser436, located on the cytoplasmic face of the transport domain as our internal site, with a predicted E_FRET_ change from ∼0.70 and ∼0.12 upon transition from the C_o_-C_o_to the C_i-_C_i_ state. Single VcINDY homodimer E_FRET_ *versus* time trajectories (*i*.*e*., E_FRET_ trajectories) recorded using the external and internal smFRET constructs should then report upon the same set of global VcINDY conformational dynamics involved in substrate translocation but from different perspectives.

Biochemically active variants of the two VcINDY smFRET constructs designed above were created, overexpressed, purified, and labeled with donor and acceptor fluorophores to generate the externally and internally labeled smFRET constructs (VcINDY^ext^ and VcINDY^int^, respectively) (see Materials and Methods) (Fig. S1). Both VcINDY^ext^ and VcINDY^int^ were able to catalyze substrate translocation in liposomes, and exhibited transport activities that were only three-fold reduced relative to wildtype VcINDY and were similar to the activity of the unlabeled cysteine-containing mutants (Fig. S2). To investigate the conformational dynamics of VcINDY in a membrane, we reconstituted VcINDY^ext^ and VcINDY^int^ into lipid bilayer nanodiscs (Fig. 2) (20). Nanodisc-based experimental conditions are ideal for interrogating the C_o_ to C_i_ transition at equilibrium, since both sides of the membrane are exposed to identical solutions. Furthermore, the nanodiscs used here have a diameter of ∼108 Å (21), which helped establish single-molecule resolution by ensuring that only a single VcINDY dimer could occupy each nanodisc. After chromatographic purification, these nanodiscs appeared well dispersed when imaged using negative stain electron microscopy (Fig. S3).

**Figure 2.**
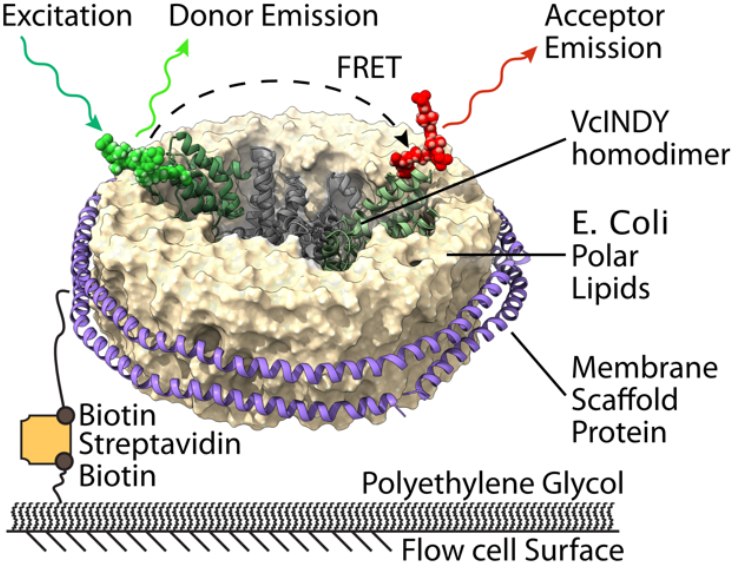
Schematic diagram of nanodisc-embedded VcINDY homodimer construct. Single cysteine mutant VcINDY labeled with a 1:1 ratio of donor and acceptor fluorophores is mixed with biotinylated membrane scaffold protein (MSP) and *E. coli* polar lipids to make nanodisc-embedded dimers. VcINDY containing nanodiscs are then tethered to a flow cell surface *via* a biotin-streptavidin-biotin bridge, and imaged using TIRF microscopy. The scaffold domains are shown in gray, and transport domains are shown in green.

smFRET experiments were performed using a widefield, total internal reflection fluorescence (TIRF) microscope to record the fluorescence intensity emitted from individual, immobilized VcINDY-containing nanodiscs (Fig. 2). The corresponding E_FRET_ trajectory for each molecule was then calculated using E_FRET_ = I_acceptor_/ (I_donor_+ I_acceptor_), where I_x_is the observed intensity of either the donor or the acceptor fluorophore. Along with the chromatographic purification used in nanodisc reconstitution, the selection criteria for E_FRET_ trajectory (see Materials and Methods), ensured that each E_FRET_ trajectory originated from a single VcINDY dimer (Fig. S1).

### E_FRET_ states consistent with elevator-type motions are stochastically sampled during VcINDY substrate translocation

To characterize the conformational dynamics of VcINDY during the transport cycle (Fig. 1B), we performed equilibrium smFRET experiments using VcINDY^ext^ or VcINDY^int^ in which the buffer contained saturating concentrations of both Na^+^ and succinate co-substrates (Buffer HS: <u>H</u>igh Na^+^ with <u>S</u>uccinate). The conformational dynamics that occur under these conditions are the same as those involved in vectorial transport—even in the absence of a chemical free energy driving force. Thus, by saturating the VcINDY smFRET constructs with both co-substrates, we expected to record E_FRET_ trajectories that report on the equilibrium conformational dynamics of the substrate translocation reaction between a substrate-bound C_o_ state and a substrate-bound C_i_ state (C_o_’’ and C_i_’’ in Fig. 1B), which we hypothesized resemble the C_o_-C_o_and C_i_-C_i_ conformations, respectively (Fig. 1C).

In the presence of both Na^+^ and succinate, the E_FRET_ trajectories originating from individual VcINDY^ext^ or VcINDY^int^ dimers exhibited stochastic, instantaneous jumps between transiently stable states characterized by distinct E_FRET_ values (*i*.*e*., E_FRET_ states) (Fig. 3). Visual inspection showed that individual E_FRET_ trajectories could visit many distinct E_FRET_ states (*e*.*g*., Fig. 3, lower panels), and the same E_FRET_ states were often revisited within a single E_FRET_ trajectory. Most E_FRET_ trajectories, however, exhibited just a few jumps to a small number of distinct E_FRET_ states before relatively rapid photobleaching of the donor and/or acceptor fluorophores prematurely ended our ability to measure E_FRET_. Qualitatively, the lifetimes of these E_FRET_ states were generally several seconds. Notably, the E_FRET_ states that are observed across the entire population of VcINDY^ext^ or VcINDY^int^ E_FRET_ trajectories (Figs. 3 and 4) are consistent with the range of predicted E_FRET_ values for large, elevator-type conformational transitions between the C_o_-C_o_and C_i_-C_i_ conformations using the donor-acceptor pairs engineered into both VcINDY^ext^ and VcINDY^int^ (Fig. 1C).

**Figure 3.**
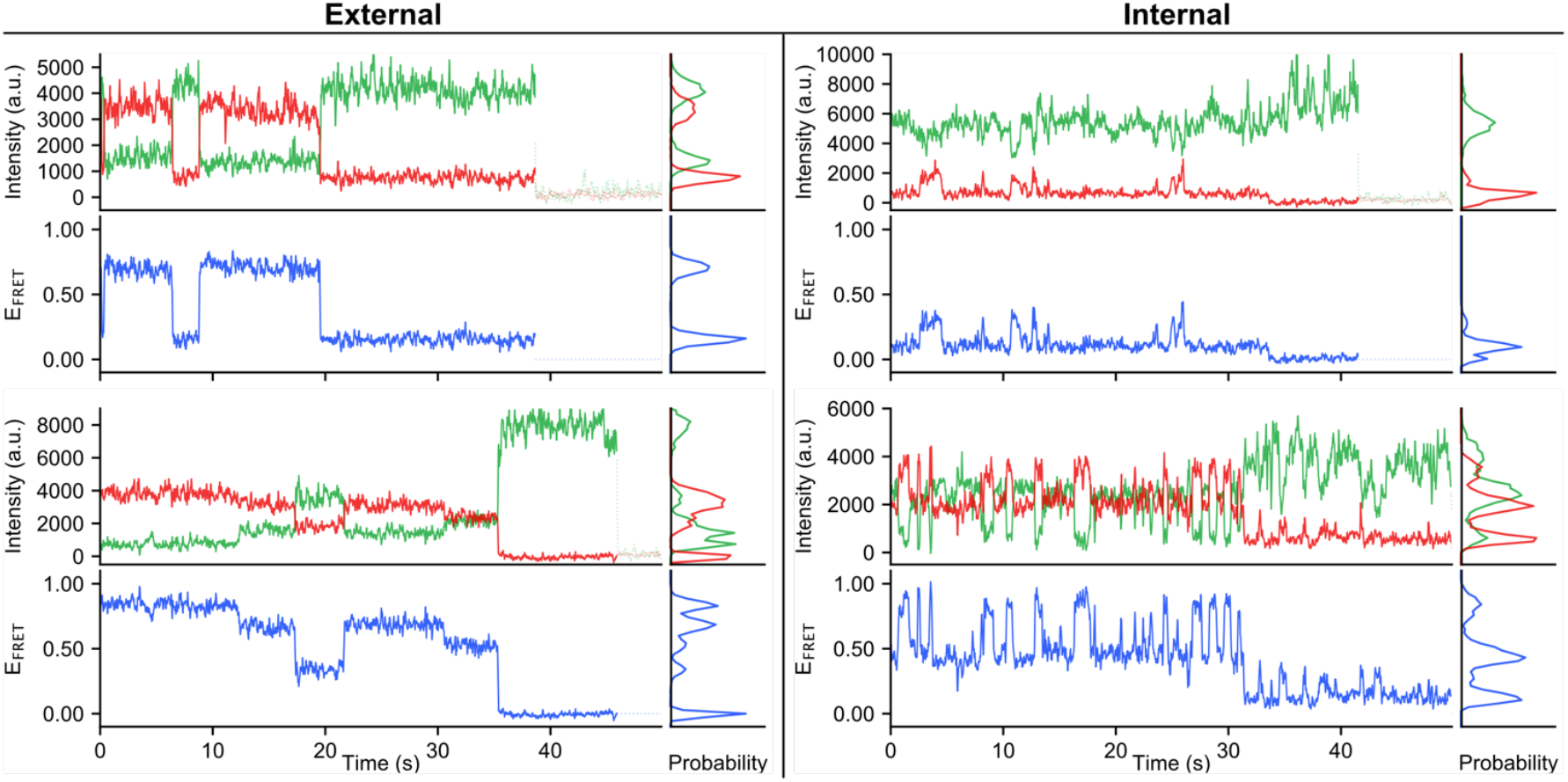
Equilibrium smFRET from VcINDY^ext^ (left) and VcINDY^int^ (right) in the presence of sodium and succinate (buffer HS). Example donor (green) and acceptor (red) fluorescence intensity trajectories, with corresponding E_FRET_ trajectories (blue) and idealized trajectories (black) are shown. Several trajectories shown include donor fluorescence intensity after the acceptor fluorophore has photobleached, but this was removed prior to further analysis.

**Figure 4.**
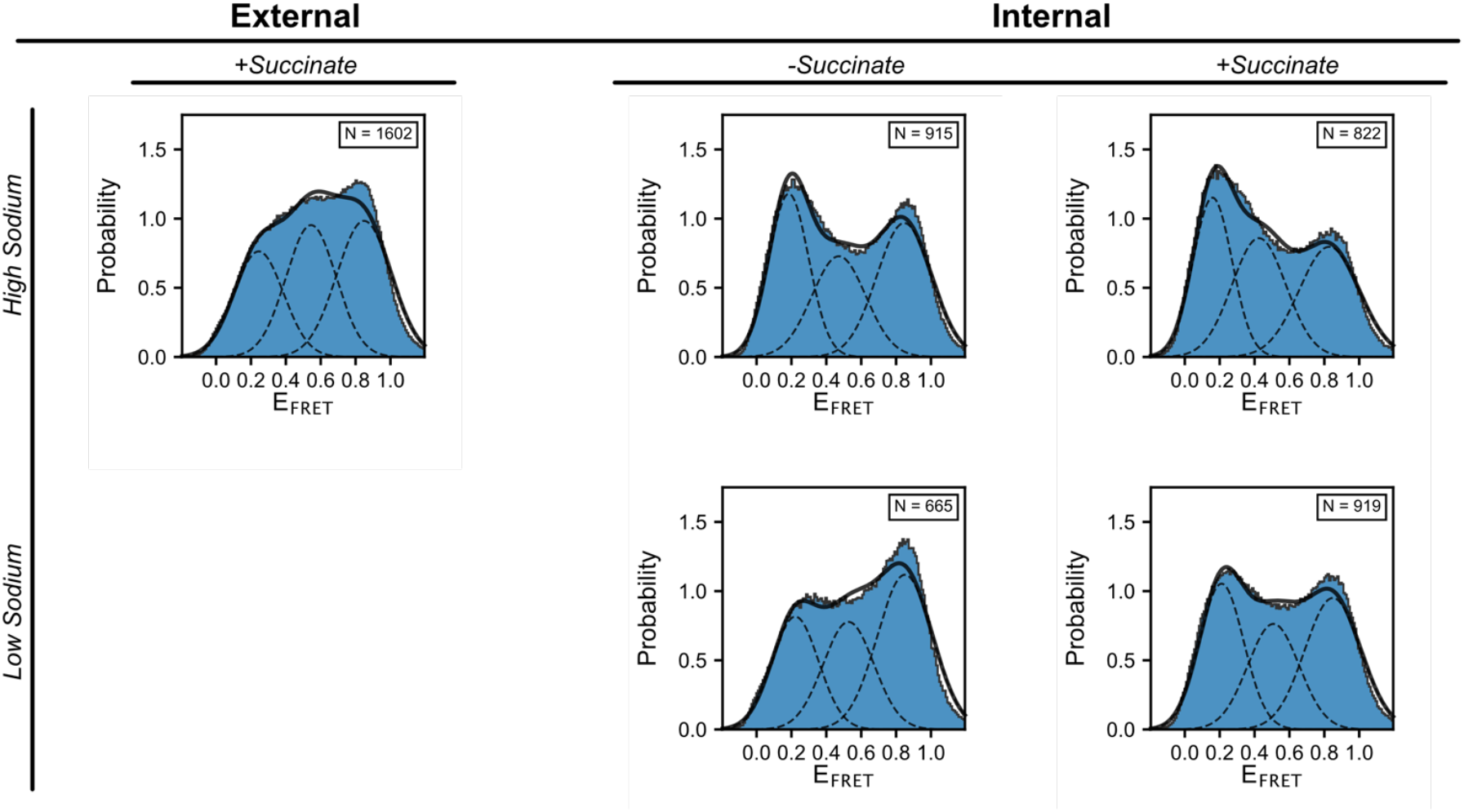
Substrate dependence of VcINDY E_FRET_ distributions. Histograms of E_FRET_ values from all external (left) and internal (right) smFRET signals recorded with VcINDY^ext^, and VcINDY^int^, respectively, in different buffer conditions. *N* refers to the number of individual single-molecule E_FRET_ trajectories contributing to each histogram and used in subsequent analyses. The E_FRET_ data in these histograms were not filtered, and E_FRET_ values corresponding to donor photobleaching have been removed (*i*.*e*., the last dwell in an E_FRET_ state of zero). The dashed black curves in each histogram denote the population distributions of the three states that are obtained from kinetic modeling, while the solid black curve denotes the total population distribution from all three states of the kinetic model.

### VcINDY reversibly populates C_o_-C_o_, C_i_-C_i_, and mixed C_o_-C_i_ conformational states during substrate transport

Assuming transmembrane substrate translocation requires transitions solely between the C_o_-C_o_and C_i_-C_i_ conformations of VcINDY, we predicted that E_FRET_ trajectories from both VcINDY^ext^ and VcINDY^int^ should exhibit transitions between only two E_FRET_ states (Fig. 1C). Instead, we observed transitions between more than two E_FRET_ states for both constructs (Figs. 3-4). The simplest interpretation of this observation is that the two protomers within a VcINDY dimer can independently undergo elevator-type conformational transitions. If that were the case, VcINDY dimers could then sample additional, ‘mixed’ conformations (*e*.*g*., C_o_-C_i_). Although none of the structural studies of VcINDY published to date support the existence of such mixed conformations, other dimeric secondary active transporters with the same fold as VcINDY have been observed to populate a C_o_-C_i_conformation (22–24). To determine whether our smFRET data are consistent with such an interpretation, we used a state-of-the-art smFRET data analysis algorithm implemented in the tMAVEN software package to computationally determine the most parsimonious kinetic model that captures the broad range of E_FRET_ states and transition dynamics observed across the entire set of E_FRET_ trajectories obtained from each construct, VcINDY^ext^ and VcINDY^int^ (25) (see Materials and Methods) (Figs. S4-S5).

For both VcINDY^ext^ and VcINDY^int^, the most parsimonious kinetic model that can simultaneously account for the entire set of E_FRET_ trajectories was a three-state model (Figs. 4, S6-S7). Notably, this was also true for smFRET data recorded under the four different combinations of sodium and succinate concentrations we investigated using VcINDY^int^ (discussed below). Because these three states persisted across different smFRET constructs and different sodium and succinate concentrations (Figs. 4, S6-S7), we concluded that our smFRET data primarily report on transitions between three distinct conformations of the VcINDY dimer. Nonetheless, individual E_FRET_ trajectories sampled more than three E_FRET_ states that, given the high signal-to-background ratio of the data, were unambiguously distinct from one another (Fig. 3). This observation suggests that each of the three states in our kinetic model is actually composed of a collection of VcINDY conformations that, although globally similar, exhibit enough local structural variation near one or both of our fluorophore-labeling sites so as to give rise to distinct E_FRET_ states.

Based on the alternating access principle, we expected that the C_o_ and C_i_ conformations of the transport cycle (Fig. 1B) should be represented within the three states of our kinetic model (2) (Fig. 4). For VcINDY^ext^, the state characterized by an average E_FRET_ value of 0.25 is consistent with our prediction of E_FRET_ = 0.20 for the C_o_-C_o_conformation, and the state characterized by an average E_FRET_ value of 0.85 is consistent with our prediction of E_FRET_ = 0.93 for the C_i_-C_i_ conformation (Figs. 1C, S7). Likewise, for VcINDY^int^, the state characterized by an average E_FRET_ value of 0.83 is consistent with our prediction of E_FRET_ = 0.70 for the C_o_-C_o_ conformation, and the state characterized by an average E_FRET_ value of 0.16 is consistent with our prediction of E_FRET_ = 0.12 for the C_i_-C_i_ conformation (Figs. 1C, S7). Thus, the conformations of VcINDY that are sampled in our smFRET data are broadly consistent with the elevator mechanism suggested by previous structural work and therefore illustrate how stochastic VcINDY conformational changes are coupled to its transport cycle.

There are many effects that could give rise to the small differences we see between our observed and predicted E_FRET_ values (ΔE_FRET_ ∼ 0.1). These include (*i*) differences between the C_o_-C_o_and/or C_i_-C_i_ conformations that nanodisc-embedded VcINDY adopts in our smFRET experiments and the corresponding conformations that are observed in the X-ray, cryo-EM, and computational studies (10, 13, 14, 17); (*ii*) structural perturbations of VcINDY due to fluorophore labeling; (*iii*) unaccounted-for contributions the linkers used to covalently attach the fluorophores to the protein make to the distance between fluorophores; (*iv*) constraints on fluorophore mobility due to interactions with the protein and/or membrane; or even (*iv*) photophysical complications such as ‘photoblinking’ of the fluorophores. Nonetheless, given the consistency of our experimental E_FRET_ values with the predicted E_FRET_ values for the C_o_-C_o_and C_i_-C_i_ conformations, we were able to assign two of the states in our three-state kinetic model to specific VcINDY conformations involved in substrate translocation. Specifically, we assigned the state in our kinetic model that exhibited average E_FRET_ values of 0.25 and 0.83 for VcINDY^ext^ and VcINDY^int^, respectively, to the C_o_-C_o_ conformation. Likewise, we assigned the state in our kinetic model that exhibited average E_FRET_ values of 0.85 and 0.16 for VcINDY^ext^ and VcINDY^int^, respectively, to the C_i_-C_i_ conformation. Notably, these assignments are self-consistent in that the most populated VcINDY conformation is the C_i_-C_i_ conformation, regardless of whether it is the VcINDY^ext^ or VcINDY^int^ smFRET data that are analyzed (Fig. 4).

The remaining, unassigned state in our kinetic model can be explained as VcINDY in either a mixed C_o_-C_i_ conformation, or a unique intermediate conformation that is distinct from C_o_-C_o_ and C_i_-C_i_. Based on biochemical and structural considerations, we favor assigning this state to the more parsimonious, mixed C_o_-C_i_ conformation. The existence of such a conformation is only possible if the protomers of VcINDY operate independently—a possibility that is consistent with the results of previous VcINDY succinate transport kinetic measurements (8). Additionally, the two-fold rotational symmetry of the VcINDY dimer renders the C_o_-C_i_ and C_i_-C_o_ conformations structurally degenerate such that they would be expected to correspond to a single E_FRET_ value, which is consistent with our results. In contrast, similar symmetry considerations for VcINDY with a unique intermediate protomer conformation would yield up to six distinct VcINDY dimer conformations in the absence of strict protomer cooperativity or anti-cooperativity. Finally, computational alignment of the VcINDY scaffold domain of the C_o_-C_o_ and C_i_-C_i_ conformations shows that the donor and acceptor fluorophores in a mixed C_o_-C_i_ conformation would be ∼50 Å apart for VcINDY^ext^ and ∼59 Å apart for VcINDY^int^—distances that yield predicted E_FRET_ values of 0.53 and 0.29, respectively. Given the caveats described above, these predicted E_FRET_ values are in reasonable agreement with the average E_FRET_ values of 0.54 and 0.42 for VcINDY^ext^ and VcINDY^int^, respectively, that characterize the remaining state of our three-state kinetic model (Figs. 4, S6-S7).

Regardless of whether the VcINDY^ext^ or VcINDY^int^ constructs are used, our kinetic modeling shows that the elevator motion of a single protomer in the VcINDY dimer translocating substrates through the membrane occurs at a rate of ∼0.4 s^-1^ (*i*.*e*., once every ∼2.5 seconds) with no apparent directional dependence. Notably, direct transitions between C_o_-C_o_ and C_i_-C_i_ where both protomers undergo an apparently simultaneous transition within the temporal resolution of our TIRF microscope were nearly an order of magnitude slower (∼0.04 s^-1^), and therefore much less likely to occur. This suggests that the individual protomers of a VcINDY dimer primarily undergo the translocation reaction in an independent and relatively uncoordinated manner. Overall, in these experiments, the elevator motions of VcINDY responsible for substrate translocation are approximately an order of magnitude faster than the turn-over rate for transport measured in bulk biochemical transport experiments of ∼0.02 s^-1^ (8). Along with the fact that our smFRET experiments were performed at lower a temperature (21 °C *versus* 30 °C), this suggests that VcINDY undergoes multiple substrate translocation reactions before a productive transport event occurs, and that the rate-limiting step of VcINDY, and possibly DASS family, catalyzed transport may involve substrate binding and/or dissociation.

### VcINDY exhibits conformational dynamics that are largely substrate independent

The basic mechanistic model of a transport cycle for VcINDY makes several fundamental predictions regarding the conformational dynamics that occur during transport (Fig. 1B). Consistent with the alternating access hypothesis (2), VcINDY should be able to switch from a C_o_ to a C_i_ conformation in the *apo*, substrate-free state. Also, to preserve substrate coupling, we expect that the partially bound transporter (C_o_’ and C_i_’ in Fig. 5) will not undergo outward to inward transitions (and vice versa) to minimize uncoupled slippage of Na^+^ or succinate. To test these predictions, we performed additional smFRET experiments using VcINDY^int^ in the absence of both co-substrates (*i*.*e*., in the *apo* condition) (Buffer LX: <u>L</u>ow Na^+^ with <u>No</u> succinate) as well as in the presence of high concentrations of one co-substrate, but the absence of the other (*i*.*e*., in the only Na^+^-bound or the only succinate-bound conditions) (Buffer HX: <u>H</u>igh Na^+^ with <u>No</u> succinate; Buffer LS: <u>L</u>ow Na^+^ with <u>S</u>uccinate). Because of the consistency we observed between VcINDY^int^ and VcINDY^ext^ in buffer HS (Figs. 3,4), these substrate-dependence experiments were performed using only the VcINDY^int^ construct. Where appropriate, we replaced Na^+^ with choline to maintain ionic strength and osmolarity. We note that all of our buffers likely contain trace (*i*.*e*., ‘low’) amounts of contaminating Na^+^, but at several orders of magnitude lower than the Michaelis constant, K_M_, measured for transport (K_m_ = 41.7 mM) (8).

**Figure 5.**
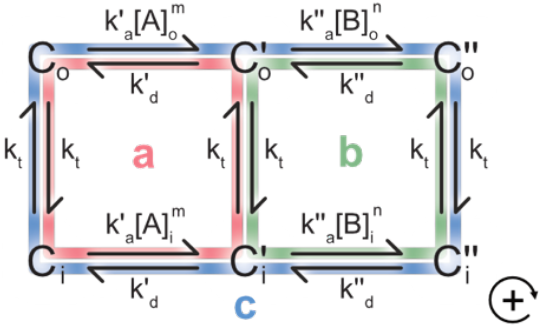
Free energy transduction model for a secondary active transporter. First order and pseudo-first order rate constants for the parsimonious kinetic model discussed in the text are given over each transition. For simplicity, co-substrate binding has been reduced to a strictly sequential, two-step process with mirror symmetry (1). Substrate A is the first co-substrate to bind with stoichiometry *m*; co-substrate B binds second with stoichiometry *n*. Prime signifies the first co-substrate is bound; double-prime signifies both co-substrates are bound; C_o_ and C_i_ denote the outward and inward facing conformation of the transporter. *k*_*t*_ is the translocation rate constant, *k*_*a*_is a second order association rate constant, and *k*_*d*_ is a dissociation rate constant. Cycles *a* (red) and *b* (green) are responsible for slippage, while cycle *c* (blue) is responsible for transport. The positive direction is defined to be clockwise.

As observed for the smFRET experiments recorded in the presence of high concentrations of both co-substrates (Buffer HS), the smFRET experiments recorded under all three additional co-substrate conditions (Buffers LX, HX, and LS) yielded E_FRET_ trajectories that exhibited stochastic, instantaneous jumps between E_FRET_ states spanning a comparably wide range as was seen in Buffer HS (Fig. 4). Computational kinetic analysis showed that the most parsimonious kinetic model that captures the broad range of E_FRET_ states and transition dynamics observed across the entire set of E_FRET_ trajectories recorded under each set of buffer conditions was again a three-state model (Fig. S6). Notably, the E_FRET_ values and transition rate constants obtained under each of these three buffer conditions were similar to each other and to those obtained under the Buffer HS condition (Fig. S7). While there were slight differences in the equilibrium population distribution of the three states observed in each buffer condition, the kinetics were relatively unchanged, with single protomer translocation reactions occurring at a rate of ∼0.5 s^-1^ under all buffer conditions tested. Thus, our quantitative kinetic analysis reveals that the conformational motions of VcINDY that are probed along the coordinate between the donor and acceptor fluorophore in VcINDY^int^ are relatively independent of the absence or presence of saturating concentrations of either or both co-substrates. Although these results conform to our expectation for the *apo* state, the observation of frequent transitions in the E_FRET_ trajectories recorded under the single-co-substrate conditions was unexpected.

### Thermodynamic modeling shows that compensatory tuning of co-substrate binding and dissociation can minimize slippage cycles

In search of an explanation for the surprising kinetics observed under the single-co-substrate conditions, we next analyzed the thermodynamic efficiency of the VcINDY transport cycle. We modeled the free-energy transduction of a secondary active transporter using Hill’s method of cycles (26) to determine how the different molecular processes involved in transport might contribute to slippage. For simplicity, we considered co-substrate binding as a strictly sequential, two-step process with mirror symmetry (*i*.*e*., the same order of co-substrate binding and dissociation on both faces of the membrane). Specifically, we considered the most parsimonious molecular interpretation of this kinetic scheme for secondary active transport, where (*i*) the rate constants are all first-order or pseudo-first order, (*ii*) the rate constants are independent of the C_o_ or C_i_ state, and (*iii*) the translocation reaction is independent of substrate binding status (Fig. 5). The resulting, simplified kinetic scheme has three Hill cycles—two of which are involved in slippage (cycles *a* and *b*) and one that is responsible for transport (cycle *c*) (Fig. 5).

We then calculated the thermodynamic efficiency, *η*, of a transporter with these assumptions (Fig. 5) (26). For a maximally efficient transporter, *η ≈*1 and the transporter tightly couples free energy from the driving to the driven co-substrate with very little slippage. In that situation, the thermodynamic efficiency is *η*∝1 − (*η*_*ac*_ − *η*_*bc*_) +⋯with small higher-order terms; here we have assumed *A* is the driving substrate, but the conclusions discussed below are independent of this. Under our assumptions,

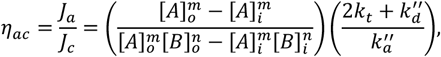

And

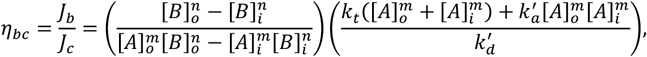

where *J*_κ_ is the cycle flux for cycle κ; concentrations are indicated by square brackets; *A* and *B* are the first and second binding co-substrates with stoichiometries *m* and *n*, respectively; the translocation rate constant is *k*_*t*_, association and dissociation rate constants are *k*_*a*_ and *k*_*d*_, respectively; and prime and double-prime denote the first and second binding reactions, respectively (Fig. 5). The first terms in these equations show that the relative amount of slippage that a transporter undergoes depends on the instantaneous thermodynamic driving forces *via* the co-substrate chemical gradients, while the second terms show that it also depends on the translocation and substrate binding and dissociation reaction kinetics. Thus, the only reliable way nature can optimize this transporter into a highly efficient free energy transducer over a broad range of environmental conditions is to have evolved control over slippage *via* evolutionary tuning of these rate constants to reduce the total magnitude of *η*_*ac*_ and *η*_*bc*_. This suggests that optimal transporters have faced evolutionary pressure to decrease 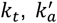 and/or 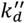,and to increase 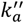 and/or 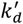 such that the first binding co-substrate weakly binds until the second arrives and then both co-substrates can tightly bind. Unfortunately, tuning these rate constants will also affect the overall transport cycle flux, *J*_*c*_, and this creates a speed-accuracy tradeoff.

The speed-accuracy tradeoff can, however, be navigated. At the molecular level, the clearest way to reduce slippage is for the binding pocket to stabilize the *apo* and fully bound states (C and C”), by some amount Δ*G*^′^ and Δ*G*^′′^, respectively, and to source all of that energy from a compensatory destabilization of the partially bound conformation, C’, by an amount Δ*G*^′^ + Δ*G*^′^′, all without affecting the transition state energy barriers (Fig. 6). This is a conceptually similar idea to the use of substrate binding energy to stabilize the transition state for enzyme-substrate transition state complementarity (27). In our case, this isoenergetic change increases 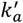 and 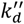 while decreasing 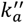 and 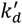,which reduces slippage and increases the thermodynamic efficiency of the transporter. It also has the effect of increasing the transport cycle flux *via* ∼75% of the terms defining *J*_*c*_ (see Supporting Information for the expression for *J*_*c*_). Given this substantial ability to speed up transport in the complicated expression for *J*_*c*_, we suspect that these compensatory, isoenergetic binding and dissociation reaction tunings optimally navigate the speed-accuracy tradeoff by simultaneously decreasing the relative fraction of slippage while increasing the total amount of transport. The exact extent of these abilities, however, depends on the instantaneous internal and external co-substrate concentrations, which can vary significantly across physiologically relevant conditions (Fig. S8) (28). Regardless, this thermodynamic modeling of the free energy transduction efficiency of a secondary active transporter predicts that slippage can be reduced simply by isoenergetic optimization of the energy landscape of the co-substrate binding reaction (Fig. 6). In other words, increased cooperativity of co-substrate binding can itself lead to minimized slippage.

**Figure 6.**
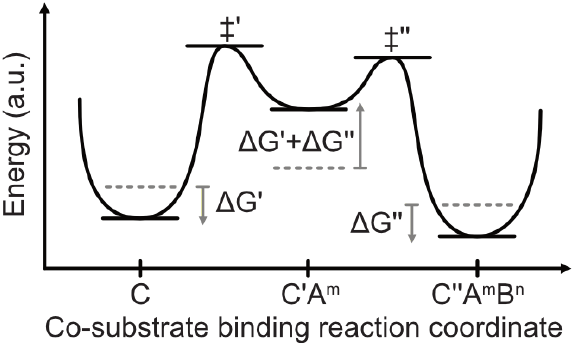
Schematic of optimized energy landscape for binding co-substrates A and B. Original energy levels are denoted with dashed lines for apo-(*C*), partially bound (*C*′), and fully bound (*C*′′) transporter. Direction of perturbations to energy levels by amounts Δ*G*^′^ and Δ*G*′′ are shown with arrows. Relative positioning of unperturbed *C, C*′, and *C*′′, and transition states ‡ ′ and ‡ ′′ are arbitrarily chosen, as are the magnitudes of Δ*G*^′^ and Δ*G*′′.

Finally, we also used this thermodynamic modeling-based approach to estimate the occupancy of *apo* VcINDY in our smFRET experiments performed in the presence of only saturating Na^+^ (Buffer HX). Specifically, we modeled the reported *K*_*M*_ and *V*_*max*_values for the Na^+^ titration and succinate titration experiments that recorded succinate uptake rates in the biochemical proteoliposome-based transport assay data of Mulligan and coworkers (see Supporting Information for extended details) (8). This was an underdetermined problem, so we assumed that a single, average rate constant defined several molecular processes: 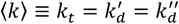,Markov chain Monte Carlo optimization yielded estimates of 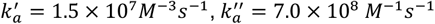,and ⟨*k*⟩ = 1.05 × 10^3^ *s*^−1^ (Figs. S9-S10), and reasonably matched the original titration data (Fig. S11). The small mismatch is due to the assumption of a single, average rate constant and the uncertainty in the binding order of the three Na^+^ ion- and succinate co-substrates. Regardless, these estimates are consistent with the biochemical transport data in the context of our kinetic mechanism, so we used them to calculate the steady-state probability of *apo* VcINDY in Buffer HX to be 6.5% (see Supporting Information). This estimate suggests that even with 100 mM Na^+^ in solution, VcINDY still spends a non-negligible portion of time with an unoccupied binding pocket. Given this result, it is highly likely that the majority of the unexpected elevator motions present in the E_FRET_ trajectories recorded in the presence of only one co-substrate originate from *apo*, rather than partially bound, VcINDY.

### The dicarboxylate substrate binding pocket of VcINDY is enthalpically stabilized by sodium ions

To obtain atomic-resolution insight into the Na^+^ and dicarboxylate substrate binding reaction for the VcINDY transport cycle, we determined how the conformational state of the VcINDY homodimer is influenced by different dicarboxylate substrates. We therefore separately crystallized detergent-solubilized VcINDY with Na^+^ and three different C_4_-dicarboxylate substrates (succinate, fumarate, and malate) and solved their structures at 3.09 Å, 3.29 Å, and 3.50 Å, respectively. Regardless of the dicarboxylate substrate identity, VcINDY adopted the same C_i_-C_i_
state in all three of these structures. Similar to what was seen in the previous structure of the C_i_-C_i_ state (13, 14, 17, 18), the binding pocket of the dicarboxylate substrate in all three of these new structures was partially comprised of two hairpin loops that coordinated distinct Na^+^ ions, Na1 and Na2 (Figs. 7, S12). Given this binding geometry, it is possible that tight binding of both co-substrates requires stabilization of these hairpin loops and that their local conformational rearrangements gate binding and/or dissociation of the co-substrates.

**Figure 7.**
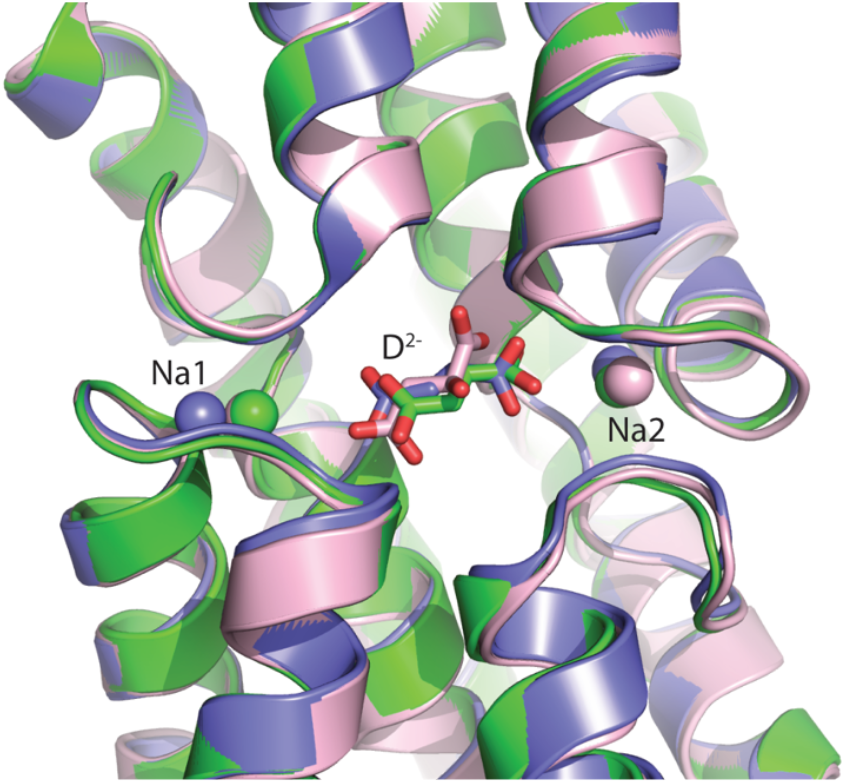
View of the VcINDY binding pocket from the cytosol. The X-ray crystallography derived structural model is of dicarboxylate (D^2-^) and Na^+^-bound VcINDY for malate (pink), succinate (blue), and fumerate (green).

## Discussion

### The VcINDY transport domain undergoes elevator motions to enable alternating access

The ubiquitous presence in our experiments of stochastic, instantaneous transitions between E_FRET_ states consistent with the C_o_-C_o_ and the C_i_-C_i_ conformations is the first direct evidence for the hypothesis that VcINDY uses an elevator transport mechanism to enable alternating access (Fig. 3) (10). Because we directly observed these stochastic, elevator motions in the presence of substrates but absence of an electrochemical gradient (Buffer HS) and also in the absence of any co-substrates at all (Buffer LX), it is clear that the VcINDY transport domain can translocate the substrate binding site across the membrane using only thermal energy—as is expected for the conformational motions of a secondary active transporter that are responsible for transport. Moreover, the rate constants for that process are on the order of 10^−1^ s^-1^ (Fig. S7), suggesting that the energy barrier separating the C_o_ and C_i_ states along the translocation reaction coordinate is not prohibitively high. Since the VcINDY transport cycle is much slower and occurs with a rate on the order of 10^−2^ s^-1^ (8), it seems that multiple back-and-forth substrate translocation events occur before a productive transport event is completed. This observation is consistent with a model in which a molecular process other than transport domain translocation govern the transport rate (*e*.*g*., gating dynamics within the substrate binding pocket itself).

### The conformational dynamics of VcINDY protomers are non-cooperative

The experimental observation of a mixed C_o_-C_i_ conformation for the VcINDY dimer constrains the role that the two protomers can play in the transport cycle. Previously, proteoliposome-based transport experiments determined that VcINDY transports succinate with a Hill coefficient of 0.88 ± 0.13 (8). Given the VcINDY transport stoichiometry of 3:1 Na^+^:dicarboxylate (9), this Hill coefficient measurement suggested that the VcINDY transport cycle is non-cooperative and that each VcINDY protomer can independently transport succinate. Our observation of a mixed C_o_-C_i_ conformation reveals that VcINDY protomers can undergo independent substrate translocation reactions during vectorial transport, providing a structure-based mechanism for the non-cooperative transport cycle of VcINDY. Specifically, since the VcINDY dimer can significantly occupy the C_o_-C_i_ state, the two VcINDY protomers must not undergo a strictly cooperative translocation reaction (*i*.*e*., C_o_-C_o_ ⇌ C_i_-C_i_). Furthermore, they must not undergo a strictly anti-cooperative translocation reaction either (*i*.*e*., C_o_-C_i_ ⇌ C_i_-C_o_), because this behavior would preclude occupancy of the C_o_-C_o_ and C_i_-C_i_ conformations observed in our smFRET experiments and structural studies. Therefore, our results provide direct evidence that the VcINDY protomers undergo non-cooperative translocation reactions, and that the VcINDY transport cycle itself can be considered to occur independently in each protomer.

In support of this idea, recent structures of CitS, another homodimeric elevator transporter in a different structural family, reveals a CitS dimer with one inward-facing and one outward-facing protomer (22-24), and a recent high-speed atomic force microscopy study of the CitS elevator translocation motion reported that CitS undergoes non-cooperative interprotomer conformational dynamics (29). Combined with evidence that Glt_Ph_, an elevator-type, glutamate transporter that, unlike VcINDY, is from a thermophilic organism, also shows independence between the motions of its three protomers (30-32), these results suggest that such independence may be a general feature of oligomeric elevator transporters (33).

### Slippage cycles are avoided without inhibiting VcINDY transport domain translocation

To avoid wasting the free energy provided by the Na^+^ gradient that drives transport of substrates across the membrane, the conformational motions of VcINDY must be coordinated. In particular, VcINDY must minimize slippage cycles in which Na^+^ ions are transported across the membrane independent of dicarboxylate (Fig. 1). We had hypothesized that VcINDY avoids such slippage cycles by inhibiting the substrate binding pocket translocation reaction (*i*.*e*., C_o_ ⇌ C_i_) when sub-stoichiometric amounts of the co-substates are bound, and by promoting those motions only when VcINDY is either empty or fully bound. However, the smFRET experiments we recorded under single substrate conditions yielded E_FRET_ trajectories that exhibited no discernable differences in the E_FRET_ states or their rates of transitions, regardless of the presence of relatively high concentrations of either individual co-substrate in solution (Fig. S7). Given our determination that those observed transitions could reasonably correspond to *apo* VcINDY, it is quite possible that transitions between C_o_ and C_i_ in partially bound states are directly suppressed and that we were not able to observe those effects, but our smFRET experiments suggest that VcINDY also uses another, additional mechanism to minimize slippage (Fig. 1B).

There are at least two additional mechanisms that VcINDY can use to minimize slippage. One possibility is a secondary conformational change—independent of C_o_ ⇌ C_i_translocation—to ‘gate’ both binding and release of co-substrates. For instance, while VcINDY could be able to translocate substrates across the membrane if the C_o_ ⇌ C_i_transition is not completely inhibited, a slippage cycle cannot be completed without co-substrate release. Mechanistically, such gating could be achieved with an additional VcINDY conformational equilibrium that is relatively independent of the transport domain translocation reaction C_o_⇌ C_i_. Without direct structural insight into such a hypothetical equilibrium, it is attractive to think that it might involve a conformational rearrangement of the binding pocket and, in particular, the hairpin loops (*c*.*f*., Fig. 7). In this scenario, the transitions we observe in the smFRET data for partially bound VcINDY would reflect transitions between so-called ‘occluded’ states.

Another possible mechanism to minimize slippage is to ensure that co-substrate binding is extremely cooperative, such that stable binding requires both co-substrates to be present. In our smFRET experiments performed in Buffers HX and LS, this mechanism would manifest as transitions in the E_FRET_ trajectories that predominately correspond to transport domain translocation of *apo* VcINDY. This alternative scenario is consistent with the non-negligible fraction of *apo* VcINDY we estimated to be present under these experimental conditions. Furthermore, it could explain the apparent lack of substrate dependence observed relative to our smFRET experiments in Buffer LX (Fig. S7). This cooperative co-substrate binding scenario agrees with our thermodynamic modeling (Fig. 5), which predicts that an optimized transporter that has evolved to minimize slippage cycle flux should have extremely stable *apo* and fully bound states with very weak binding of the initial co-substrate(s) (Fig. 6). The extent of that optimization reflects how nature has navigated the speed-accuracy tradeoff between rapid transport and minimal slippage. This raises the possibility that some secondary active transporters undergo a non-negligible amount of slippage in order to be efficient at transporting the driven co-substrate as long as they do not significantly waste the free energy of the driving co-substrate.

Experimental evidence is also consistent with a cooperative co-substrate binding mechanism. For instance, our X-ray crystallographic structures show that Na1, Na2, and dicarboxylate substrates all interact with common hairpin loop elements (Fig. 7). That the conformation of VcINDY is effectively independent of dicarboxylate identity suggests that this fully bound conformation is very stable—as predicted in our modeling of optimal co-substrate binding. Moreover, recent experiments demonstrated that in the presence of high concentrations of only Na^+^, cysteine residues engineered near the substrate binding pocket of VcINDY are protected from modification by maleimide-derivatized polyethylene glycol (11, 16) indicating some interaction between Na^+^ and the *apo* VcINDY binding pocket. This result is consistent with a Na^+^-first binding order and with predictions that slippage is minimized and that transport is optimized by weak binding of the first co-substrate(s) (Fig. 6). In future work, determination of the affinities and binding order of co-substrates, together with structure characterization of the sites, will be required to develop a comprehensive understanding of how co-substrate binding is employed as a mechanism to avoid slippage cycles during vectorial transport by VcINDY.

## Conclusion

Previous studies of the structure, transport biochemistry, and alternating access mechanism of VcINDY have established it as a prototypical membrane-bound transporter of the DASS family (13, 8, 10, 14, 9, 17, 11). Here, we have advanced this fundamental understanding of the DASS family and secondary active transport by directly determining how the conformational dynamics of VcINDY enable it to achieve alternating access and how those dynamic motions are tuned in order to maintain the efficiency of the transport cycle. Specifically, our study provides direct evidence that (*i*) the VcINDY translocation reaction uses a thermally driven, elevator mechanism; (*ii*) VcINDY protomer conformational dynamics are non-cooperative; (*iii*) the VcINDY transport cycle uses cooperative substrate binding to minimize slippage; and (*iv*) cooperative substrate binding is a general approach to optimize transport efficiency for certain secondary active transporters. Altogether, these findings immediately suggest future studies aimed at determining the extent to which highly cooperative co-substrate binding and/or additional conformational dynamics that gate transport (1, 4, 34) enable VcINDY to avoid slippage cycles during vectorial transport. Overall, the powerful smFRET-based experimental modality for interrogating the function and regulation of VcINDY established here is easily extended to study the transport cycles of other DASS family transporters, and even dimeric transporters from other families, to further our understanding of secondary active transport.

## Materials and Methods

### Expression and purification of VcINDY

Expression and purification of wild type VcINDY was carried out according to our previous protocol (13). Briefly, *Escherichia coli* (*E. coli*) BL21-AI cells (Invitrogen) were transformed with a modified pET vector encoding N-terminal 10X His tagged VcINDY. Cells were grown for 16 hours post-induction at 19°C. After cell breakage, membranes were solubilized in 1.2% N-decyl-β-maltoside (DM) and the protein was purified on a Ni-NTA affinity column (Qiagen). Following His-tag removal, VcINDY was further purified by size-exclusion chromatography (SEC) in buffer containing 25 mM Tris pH 8, 100 mM NaCl, 5% glycerol, 0.1% DM unless otherwise indicated.

### Purification of VcINDY mutants

Expression and purification of VcINDY mutants G211C and S436C was performed as with the wildtype protein, but with the following modifications. All buffers contained 10 mM sodium citrate but not glycerol. Membranes were solubilized in 1.0% N-dodecyl-β-maltoside (DDM). The protein was eluted from Ni-NTA agarose beads using 300 mM imidazole. Finally, the SEC buffer contained 25 mM Tris pH 8, 100 mM NaCl, 0.1% DDM and 0.5 mM TCEP.

### Expression, purification and biotinylation of membrane scaffolding protein

Expression and purification of membrane scaffolding protein (MSP1E3D1) was carried out according to a previously published protocol (35). In brief, *E. coli* BL21-Gold (DE3) cells (Agilent) were transformed with a modified pET 28a vector encoding N-terminal 7X His tagged MSP with a mutated cysteine at position 277 for biotinylation. Cells were grown at 30°C for 4 hrs post 0.1 mM IPTG induction at OD 0.8. After cell breakage, lysate was bound to Ni-NTA agarose beads (Qiagen) for 4 hours with nutation at 4°C. Protein was eluted with 100 - 250 mM imidazole and dialyzed into 100 mM Tris pH 7.4, 300 mM NaCl for 1 hr at 4°C. The His-tag was then removed by overnight TEV digestion at 12°C. Cleaved MSP protein was subsequently biotinylated using EZ-Link Maleimide-PEG11-Biotin (Promega) according to the manufacturer’s instructions. Specifically, a 20-fold molar excess of biotin reagent dissolved in DMSO was incubated with protein overnight at 4°C. Labeled MSP1E3D1-T277C protein was further purified by SEC in 100 mM NaCl, 1 mM TCEP, 20 mM Tris pH 7.4.

### Fluorophore Labeling

Stock solutions of Alexa Fluor 555 C2 maleimide (AF555) and Alexa Fluor 647 C2 maleimide (AF647) were prepared in anhydrous DMSO at 3mM and stored at -20°C. Prior to labeling, a 1 mg/ml solution of VcINDY (total of 0.5mg to 1mg) was treated with 10mM DTT for 2hrs at 4°C to reduce the cysteines.

DTT was then removed by 3 serial dilutions in SEC buffer (25 mM tris pH 8.0, 100 mM NaCl, 5% glycerol, 0.15% DDM, 1 mM TCEP) followed by re-concentration with an Amicon concentrator (cutoff 50 kDa). This preparation was incubated with a 10-fold molar excess of AF555 and AF647 overnight at 4°C, protected from light. The resulting solution was purified on Superdex 200 10/300 GL size exclusion column to remove the unreacted dye. Fractions from the elution peak were combined and the labeling stoichiometry was determined with a spectrophotometer using the following wavelengths and extinction coefficients: VcINDY: λ=280 nm with J = 57075 cm^−1^M^−1^; AF555: λ= 556 nm with J= 155,000 cm^−1^M^−1^; AF647: λ= 651 nm with J= 270,000 cm^−1^M^−1^. Typical labeling efficiency was 42% for each dye.

### Nanodisc Reconstitution

A thin film of *E. coli* polar lipid (EPL) was prepared by adding 10 mg of chloroform stock solution to a glass test tube. The solvent was evaporated with argon gas and the test tube was placed in a vacuum chamber at ∼1 mTorr for ∼2 hr. This lipid film was resuspended by adding 285 μl of nanodisc buffer (20mM Tris pH7.4; 100mM NaCl, 0.5mM EDTA, 0.5mM TCEP) followed by vortexing and sonication in an ultrasonic bath (Laboratory Supplies Co. Inc, Model G112SP1T) with 10 sec bursts alternating with 10 sec rests for a total of 3min. This aqueous lipid solution was solubilized by adding 215 μl of 10% DDM to produce 500 μl of final lipid stock concentration at 20mg/ml.

To reconstitute VcINDY into nanodiscs, detergent-solubilized EPL was added to VcINDY at a molar ratio of 280 and incubated on ice for 10 min. Biotinylated MSP was then added to this mixture at an 8-fold molar excess to produce 500 μl of a solution containing 2.5mM EPL, 72 μM MSP, 9 μM VcINDY and 7.2 mM DDM. This solution was incubated at 4°C for 1hr followed by the addition of SM-2 Bio-Beads in three steps: 300 mg initially, 200 mg added 60 min later, and 100 mg added at 90 min later and incubated overnight. During these incubations, solutions were kept at 4°C and gently stirred.

Nanodiscs containing VcINDY were purified by incubating the solution with 0.5 ml of Ni-NTA beads pre-equilibrated with nanodisc buffer (20 mM Tris pH=7.4; 100 mM NaCl; 0.5 mM EDTA; 0.5 mM TCEP) for 30 min at 4°C. The beads were packed into a column and washed with 1 ml of nanodisc buffer. Nanodiscs containing VcINDY were then eluted with 2 ml nanodisc buffer supplemented with 0.3 M imidazole and the peak elution fractions were pooled and concentrated with an Amicon concentrator (cutoff 50 kDa). The concentrated sample was fractionated on Superdex 200 10/300 GL size exclusion column (GE Healthcare) pre-equilibrated and eluted with nanodisc buffer. The fractions were evaluated by SDS-PAGE and those containing both MSP and VcINDY were pooled, and concentrated to ∼0.2-0.4 mg/ml. Typical yield for 1mg of VcINDY was ∼0.4 mg of reconstituted protein (VcINDY plus MSP). Electron microscopy grids were prepared in 2% (w/v) phosphotungstic acid (pH 7.5) and were examined in a Talos L120C transmission electron microscope (FEI-Thermo Fisher Scientific). Nanodisc samples were frozen in aliquots and stored at −80°C.

### Proteoliposome Reconstitution and Transport Assay

VcINDY was reconstituted into proteoliposomes as previously described (8, 10). These proteoliposomes were then used in succinate transport assays as previously described except at room temperature (8, 10). Briefly, proteoliposomes were exchanged into external buffer containing tritiated succinate with an inward facing Na^+^ gradient, samples taken at various time points were quenched and rapidly filtered over a 0.22 *μm* nitrocellulose membrane, and then succinate content inside the proteoliposomes was measured using scintillation counting. Transport rates were calculated from a linear fit to at least three data points within the first minute of the transport reaction.

### Crystallization of Wild-Type VcINDY with Substrates

Crystals of VcINDY were grown with a protein concentration of 6 mg/ml at 4°C in hanging-drop vapor diffusion. For VcINDY-succinate, crystallization was carried out with 30% PEG 300, 5 mM sodium succinate, 200 mM sodium chloride, and 100 mM MES pH 5.8. VcINDY-fumarate crystals were grown with 37.5% PEG 300, 200 μM sodium fumarate, and 100 mM sodium acetate pH 4.6. For VcINDY-malate, crystals were grown with 25% PEG 400, 20 mM sodium malate, and 100 mM sodium acetate pH 5.0.

### TIRF Microscopy

smFRET imaging experiments were performed on a prism-based, wide-field TIRF microscope at a temperature of 21 ± 1 °C. Briefly, a 532 nm diode-pumped solid-state continuous-wave laser (Laser Quantum, GEM532) was directed through an acousto-optic modulator (AOM) (Isomet, IMDD-P80L) such that the +1 order peak is directed toward a prism suspended over a 60x, 1.2 NA water immersion objective (Nikon). VcINDY embedded nanodiscs were tethered to the surface of quartz slides derivatized with dilute biotin-poly(ethylene)-glycol (PEG) in PEG *via* a biotin-streptavidin-biotin bridge, and illuminated with the TIR field generated at the slide/buffer interface. The instantaneous illumination power incident at the prism entry face was 40 mW (estimated density of 4.3 W/cm^2^), and the AOM was modulated with a 50% duty cycle at 1 MHz, such that the time averaged power was 20 mW. Fluorescence was collected through the objective, separated with a multi-wavelength imager (Photometrics, Dual-view), and imaged onto a water-cooled electron-multiplying charge coupled device (Andor, iXon Ultra 888). Movies were collected for 2000 frames at 25 msec exposures with 2x binning, 600 EM gain, 1x pre-amp gain, 30.0 MHz readout mode, 0.60 μs vertical speed, and +4 clock voltage running at −67°C using the open source microscopy software Micro-Manger (36).

### TIRF Microscopy Imaging Buffers

VcINDY homodimers were imaged in the following buffers, which all contained 100 mM KCl, and 20 mM Tris-Cl (pH_25C_=7.5). <u>Buffer LX</u>: 100 mM choline chloride. <u>Buffer LS</u>: 100 mM choline chloride, and 1 mM succinic acid. <u>Buffer HX</u>: 100 mM NaCl. <u>Buffer HS</u>: 100 mM NaCl, and 1 mM succinic acid. All buffers also contained 1% β-D-glucose, 1 mM trolox, 0.0025 U glucose oxidase, 0.02 U catalase, 1 mM 1,3,5,7-cyclooctatetraene, and 1 mM 3-nitrobenzyl alcohol as photostabilizing additives.

### TIRF Microscopy Data Analysis

Diffraction-limited spots of fluorescence intensity within the first 400 frames (10 seconds) were located in both the donor and acceptor fluorescence channels. Intensity versus time trajectories were estimated using the iterative, maximum likelihood formulas for intensity estimation 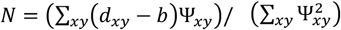 and 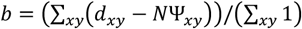, where (x,y) is a pixel location, N is the fluorescence intensity, b is the background intensity, and **Ψ** is a fixed width, Gaussian-shaped point spread function with width defined by the TIRF microscope optics (37). Donor and acceptor fluorescence channels were aligned using a 4^th^ order polynomial transform of ∼2800 control points obtained from an image of an array of sub-diffraction limit nanopatterned features localized in both channels. Trajectories were manually classified to select trajectories from only donor-acceptor labeled molecules (see Fig. S1). The photobleaching point for each E_FRET_ trajectory was inferred using Bayesian inference (see Supporting Information). A non-linear filter (38) was applied to fluorescence intensity trajectories for easier visualization in intensity and E_FRET_trajectory plots, but all analyses (*e*.*g*., generation of histograms, kinetic modeling, *etc*.) were performed using the unfiltered data.

### Kinetic Modeling of smFRET Data

Optimal kinetic models were obtained by estimating a global hidden Markov model (HMM) from all of the E_FRET_trajectories for a single experimental condition using variational Bayesian inference as implemented in the tMAVEN software (25, 39) (Fig. S4). Specifically, a variational posterior with separable emissions and kinetics was optimized using the expectation-maximization algorithm until the logarithm of the evidence lower bound (ELBO) converged to a relative change threshold of 10^−10^. For each collection of E_FRET_trajectories, model selection was performed by estimating separate HMMs with one through ten states, and then selecting the HMM with the largest ELBO (Fig. S6) as the optimal kinetic model.

## Materials, Data, and Code Availability

Mutant VcINDY and MSP expression vectors are available upon request. VcINDY structures with fumarate, malate, and succinate bound were deposited in the Protein Data Bank with IDs 6OKZ, 6OL0, and 6OL1, respectively. Code to process TIRF microscopy movies can be found in a GitHub repository (https://github.com/GonzalezBiophysicsLab/vbscope-paper) (37), and code to analyze smFRET data and kinetics is available in the tMAVEN software available in a GitHub repository (https://github.com/GonzalezBiophysicsLab/tmaven) (25).

## Acknowledgments

R.L.G. acknowledges support from the National Institutes of Health (NIH) through grants R01-GM084288 and R01-GM137608. D.N.W. acknowledges support from the NIH through grants R01-GM121994, R01-NS108151, and R01-DK099023, the G. Harold and Leila Y. Mathers Foundation, and the TESS Research Foundation. J.M. and C.M. acknowledge support from the National Institute of Neurological Disorders and Stroke (NINDS) Intramural Program, and C.M. acknowledges support from the Biotechnology and Biological Sciences Research Council through grant BB/V007424/1. D.L.S. acknowledges support from NIH through grant R35-GM144109. J.F.H. acknowledges support from NIH through grants R01-GM127883. E.T. acknowledges support from NIH through grants P41-GM104601 and R01-GM123455. D.B.S. acknowledges support, in part, from an American Cancer Society Postdoctoral Fellowship (PF-17-135-01) and from the Office of the Assistant Secretary of Defense for Health Affairs, through the Peer Reviewed Cancer Research Program under Award No. W81XH-16-1-0153. Crystal screening and X-ray data collection were carried out at NSLS-II Brookhaven National Laboratory and at Argonne National Laboratory, Structural Biology Center (SBC) at the Advanced Photon Source. SBC-CAT is operated by Univ.

Chicago Argonne, LLC, for the U.S. Department of Energy, Office of Biological and Environmental Research under contract DE-AC02-06CH11357. Electron microscopy was performed at the NYU School of Medicine Microscopy Core.

## Supporting Information for

## Supporting Information Text

### 1. Global, variational Bayes hidden Markov modeling (vbHMM)

Global vbHMMs as implemented in the tMAVEN software package (1) were estimated in a manner similar to a previously described single trajectory vbHMM implemented in the software package vbFRET (2). vbFRET is a Bayesian inference-based ‘maximum evidence’ method (3), which allowed one to determine the simplest HMM that sufficiently accounts for the complexity of the smFRET data (2, 4, 5). In this method, one can find the most parsimonious HMM for smFRET data by estimating several HMMs, each with an increasing number of hidden states, then calculating the ‘evidence’ for each HMM and choosing the one with the maximum evidence value (2). As seen in the directed acyclic graph (Fig. S4), the only difference between vbFRET, and the global vbHMM used here is that an ensemble of E_FRET_versus time trajectories is assumed to be independent and identically distributed (I.I.D.) from the same HMM parameters in the global vbHMM model. Thus, all data points then contribute to the expectation and maximization equations for the hyperparameters governing the posterior probability distribution, as opposed to only those from a single trajectory as is done for vbFRET. This is equivalent to considering one large, concatenated trajectory with vbFRET, where however there is no conditional dependence for the hidden state on the previous hidden state for neighboring data points from different trajectories. Instead, the first data point of each trajectory depends on the initial-state probability distribution denoted here by π. As can be done with vbFRET, model selection can be performed by calculating the evidence lower bound (ELBO) for each model and choosing that model with the largest value.

To demonstrate the efficacy of using model selection with global vbHMMs, we simulated multiple four-state time series, performed model selection, and compared this to an autocorrelation function (ACF) analysis (Fig. S5). The two-point autocorrelation function (6) of a stationary stochastic process or signal, y(t), describes how similar the process remains after some time lag, *τ*, and can be calculated as

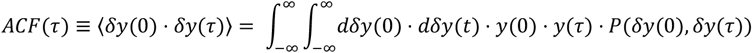

where δ *y*(*τ*) = *y*(*τ*) − ⟨*y*⟩, angular brackets denote an expectation value, P(x) is the probability distribution function of x, and

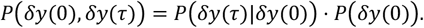

For a discrete, zero-mean signal, A(t), composed of N data points, the ACF can be estimated as

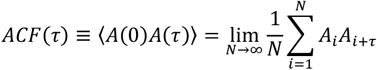

Thus, assuming normally distributed values of *A*_*i*_*A*_*i*+*τ*_, the ACF and the associated precision is distributed as a student-T distribution, *T*_*ν*_(*x*|*μ*, σ^2^) with *v* degrees of freedom, as 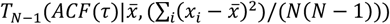,where *x*_*i*_ = *A*_*i*_*A*_*i*+*τ*_, and 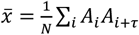.This is derived from the posterior marginal of a normal-distribution likelihood with a normal-gamma prior with *h*=0, *α*=−1/2, and *β*=0 (7). The same formulas can be used to calculate an ACF from multiple signal trajectories that are independent and identically distributed; for instance the ACF at time *τ* would then include terms 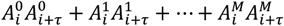,where the superscript denotes an index over *M* number of signal trajectories. We refer to this as the ensemble ACF.

For some specific types of signals, we can calculate an analytical form of the ACF for comparison to observed ACFs. First, if a time-dependent signal was stationary at zero and distributed according to *y*(*t*)*∽*𝒩(0, σ^2^), where 𝒩 denotes a normal distribution, and σ^2^ is the variance, the *ACF*(*t*) = σ^2^ ·) (*t*), where) (*t*) is a Dirac delta function, which is one when t=0, and zero otherwise. Second, consider a zero-mean Markov chain with N states that are normally distributed according to 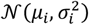 for the i^th^ state, but where ⟨*y*⟩ = 0. Also, consider that the rate constant for the transition from the i^th^ state to the j^th^ state, k_ij_, can be written as a rate constant matrix, K, with the associated Q matrix (K but where the on-diagonals are the negative sum of the rate constants out the i^th^ state). Notably, a vector of the probability of being in a particular state, P(t), given an initial probability vector, P(0), can then be written *P*(*t*) = *e*^*Qt*^ · *P*(0), where *e* denotes a matrix-exponential. If the Markov chain is in state i, the signal value at a particular time can be written *y*(*t*) *∽ μ*_*i*_ + *N*_*i*_, where 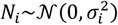.Thus, the ACF for this model is

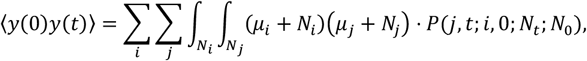

Where

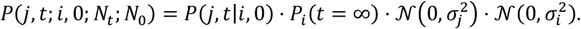

Therefore

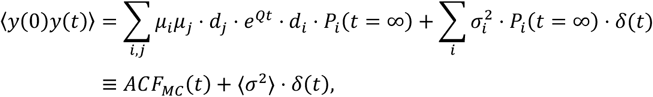

where *d*_*i*_ is a vector with a 1 in position i, and ACF_MC_is the contribution from the Markov chain. Additionally, we note that *P*(j, *t*|*i*, 0) · *P*(*i*) = *d*_j_ · *e*^*Qt*^ · *d*_*i*_ · *P*_*i*_(*t* = ∞) can be rewritten with a spectral expansion as 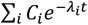,where *C*_*i*_ is a weighting factor, and λ_*i*_ is an eigenvalue of e^Qt^. Thus, for this Markov chain model with normally distributed noise, the ACF can be written as a weighted sum of *N* exponential decays with a term at t=0 corresponding to the average amount of uncorrelated, normally distributed noise. Therefore, for this model, the difference between ACF(t=0) and the next data point ACF(t=dt) is mostly ⟨σ^2^⟩, and the rest of the ACF is described by various exponential decays to zero.

In order to simulate single-molecule trajectories, we used the stochastic simulation algorithm (8) with the parameters in Table S1 with additive, normally distributed noise. From these 400 trajectories, we estimated global HMMs with one through six states as described above. As expected, the HMM with four states had the largest ELBO (Fig. S5). Additionally, we calculated the ACFs of these HMMs as described above. Notably, of these ACFs, only those for HMMs with four or more states are sufficient to match the individual and ensemble ACFs of the simulated trajectories (Fig. S5). Thus, the most parsimonious global HMM determined by this method also recapitulates the kinetic results of a model-free ACF method, but has the additional benefit as a model-based method of providing a mechanistic interpretation of those results.

### 2. Photobleaching detection

We detect photobleaching by using shape-based Bayesian model selection to choose the photobleaching time, or lack thereof, in a fluorescence intensity trajectory (9). Each model, *M*_*T*_, that we choose between is similar in that before some time, T, the data points are assumed to be distributed according to a normal distribution with some mean, μ_1_, and some precision, λ_1_(i.e., inverse variance λ=σ^−2^), which we do not care about, while the data points after time T are distributed according to a normal distribution with mean of zero, and some different precision, λ_2_. We then calculate the probability of each *M*_*T*_, having marginalized out the means and precisions so that the conclusions are relevant regardless of those values, and then use Bayesian inference to select the *M*_*T*_ that gives the best estimate of the photobleaching time as is supported by the available data. Practically, this method assigns all observed fluorescence at the beginning of a trajectory into the same ‘on’ state, usually with a large variance to account for different intensity states that compose the ‘on’ state, and then finds the last point where the data can be described as dropping to zero. For an ensemble of intensity trajectories, we perform model selection twice. The first time, we use uniform priors for the photobleaching time, while the second time, we use the independently calculated photobleaching times to infer an ensemble photobleaching rate constant, and then repeat the photobleaching time model selection calculation using this photobleaching rate constant to define the prior probability distribution for T for all of the independent intensity trajectories.

Mathematically, the likelihood for one of these models *M*_*T*_ is

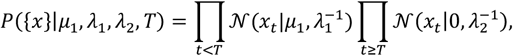

where 𝒩(*x*|*μ*, σ^2^) is a normal distribution with mean μ, and variance σ^2^. Using a prior probability distribution of

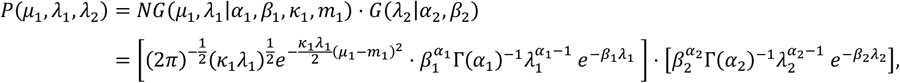

where NG is a joint normal-gamma distribution, and G is a gamma distribution,(7) we obtain a posterior probability distribution of the same form but with updated hyperparameters (denoted by primes)

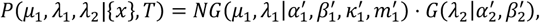

Where

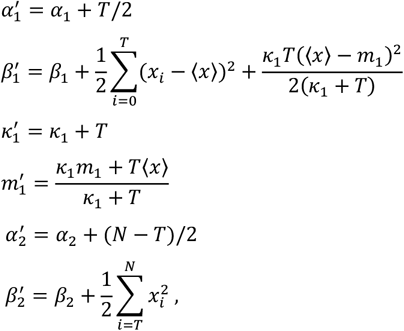

where N is the total number of data points. However, we do not particularly care about the exact values of μ_1_, λ_1_, or λ_2_, so instead, we can marginalize them out to find the probability that of the observed data given the model *M*_*T*_ (i.e., the “evidence”), regardless of these nuisance parameters. In this case, we find

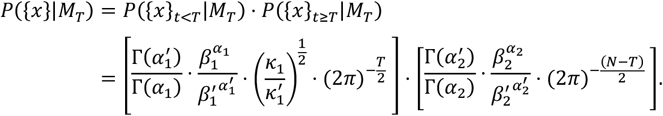

Now, with the evidence for the data given a photobleaching time, *P*({*x*}|*M*_*T*_), we use Bayesian inference to select best photobleaching model (i.e., the model best supported by the data, regardless of the individual parameters) as the largest value of

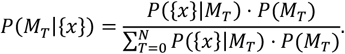

As discussed above, *P*(*M*_*T*_) is at first taken to be a constant and all trajectories are analyzed independently. Then, we infer the ensemble photobleaching rate constant using Bayesian inference with an exponential distribution as a likelihood, and a gamma distribution as the conjugate prior, and therefore also the posterior. As a result, we estimate 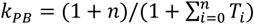,where *i* indexes the different trajectories. Then, we use an exponential distribution as the prior 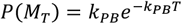,where k_PB_is treated as a deterministic value. Finally, we note that these model selection calculations are performed in log-space, and that we subtract the maximum of all the log terms (i.e., scale by a constant) from both the numerator and denominator in order the to avoid computer underflow.

Since the models used for model selection in this algorithm are not comprehensive in the sense that they do not account for every experimental nuance of the data, we believe it is best to manually examine the selected model and the underlying fluorescence intensity trajectory to ensure that they are reasonably well matched, and so that one can manually adjust the photobleaching time when they do not match due to unaccounted for experimental behaviors (*e*.*g*., deviations from Gaussian emissions).

### 3. Calculation of Cycle Fluxes

For the mechanistic scheme for transport in Fig. 5, the following terms are necessary to calculate cycle fluxes to be calculated. For shorter notation, we use 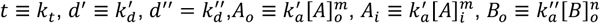,and 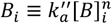.The cycle flux for any cycle is

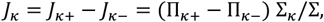

Where

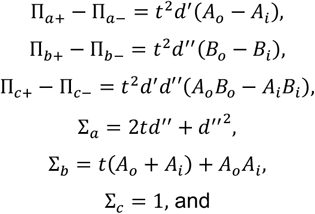

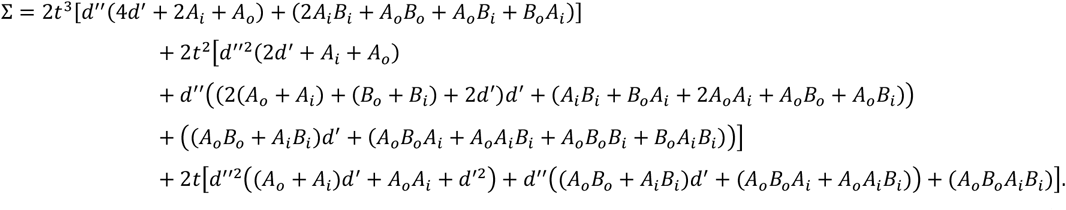

From these equations, the functional form of *J*_*c*_ for each first order rate constant, *k*, in Fig. 5 is [*ak* + *b* + *ck*^−1^]^−1^ where the coefficients *a, b*, and *c* depend on the other rate constants. This functional form has a maximum at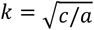,, so there is an optimal value of *k*_*t*_ that maximizes transport cycle flux under for a given set of substrate concentrations. Finally, we note that Σ has 90 terms, and thus so does *J*_*c*_. Due to the changes to the energy landscape in Fig. 6, 68 of the terms in *J*_*c*_ increase the magnitude of *J*_*c*_, 18 decrease it, and 4 do not change it.

To find the optimal parameters to describe previously published succinate transport data, a target energy function of (*J*^*expt*^ − (*J* + *J*)) ^2^was used for 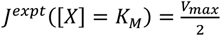 and *J*^*expt*^([*X*] = 1000*K*) = *V* for the sodium titration in their Figure 3 (*K*_*M*_ = 41.7 *mM*, and *V*_*max*_= 53.5 *kmol mg*^−1^*mik*^−1^ = 58.3 *s*^−1^), and the succinate titration in their Figure 6 (*K*_*M*_ = 1.0 *μM*, and *V*_*max*_= 232.6 *kmol mg*^−1^*mik*^−1^ = 253.4 *s*^−1^) (10). The collective energy function for both experiments was minimized using an affine invariant MCMC ensemble sampler (11) with 200 walkers, a 1,000 step burn-in period, and a 10,000 step production period from which every 100^th^ point was taken as a sample to avoid autocorrelation within the samples.

The steady-state probability of apo VcINDY in Buffer HX was calculated from the mechanism in Fig. 5 by setting [*B*]_*o*_ = [*B*]_*i*_ = 0, using the estimates for 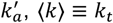,and 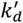 (Fig. S9) and [*A*]_*o*_ = [*A*]_*i*_ = [*Na*^+^] = 0.1*M*, to calculate the steady state probability 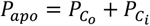 as

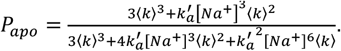

### 4. X-ray Data Collection and Structure Determination

Crystals were frozen in liquid nitrogen with the crystallization solution serving as the cryoprotectant. X-ray data were collected at the Advanced Photon Source Beamlines 19-ID and 19-BM. Data processing and scaling were performed using the HKL2000 software (12). All crystals were of space group P2_1_with unit cell dimensions around a = 106 Å, b = 102 Å, c = 171 Å, β = 99°, and contained four molecules per asymmetric unit. The structures were determined by molecular replacement using the structure of VcINDY (PDB: 5UL9) (13) with sodium and citrate removed as the initial search model, followed by repeated cycles of model building in Coot (14), and refinement in Phenix (Table S2) (15).

**Fig S1.**
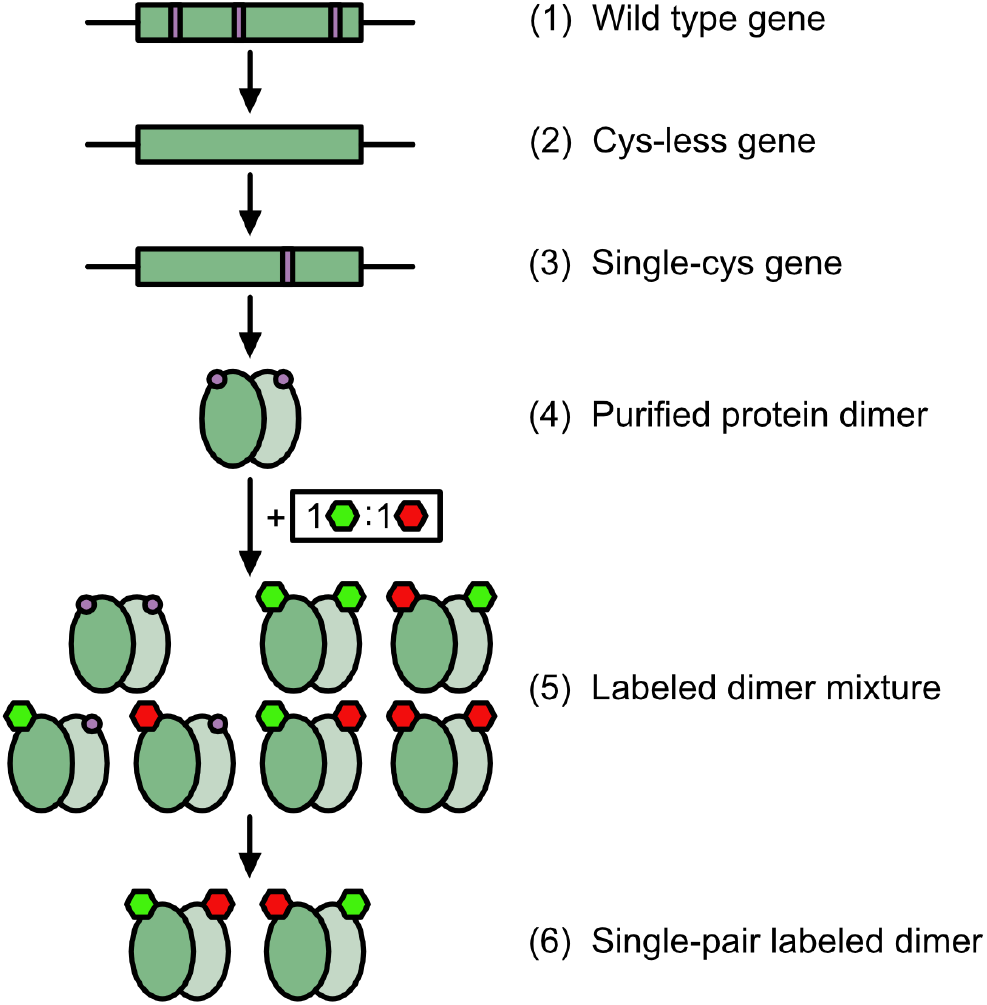
Cartoon schematic of approach to yield donor- and acceptor-fluorophore-labeled VcINDY homodimers. Site-directed mutagenesis is used to remove all codons encoding cysteine from a plasmid-based “wild type” VcINDY gene (1) to yield a “cys-less” variant (2), and used further to introduce a single codon encoding cysteine (3). This is over-expressed in *Escherichia coli* (*E. coli*), and purified to yield homodimers with single-cysteine containing protomers (4). The homodimers are labeled with a 1:1 ratio of maleimide-conjugated donor- and acceptor-fluorophores (5). Only those homodimers labeled with both a single donor- and single acceptor-fluorophore are computationally selected after imaging (6).

**Fig S2.**
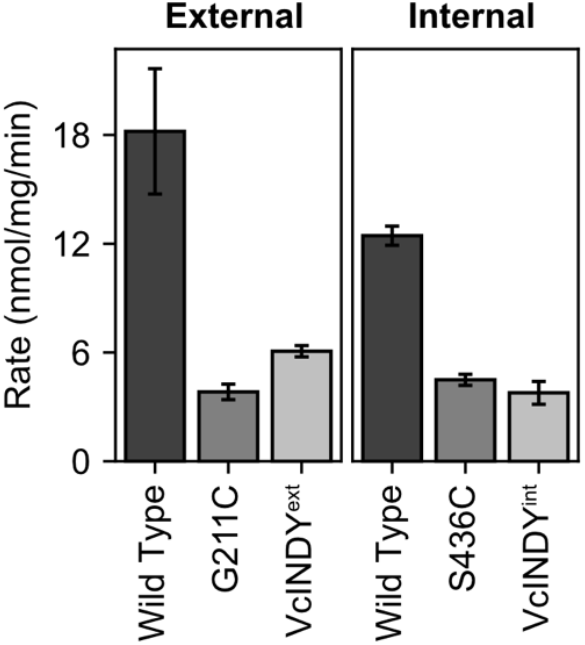
VcINDY biochemical transport assays. Transport rates of proteoliposome-embedded VcINDY homodimers for the external (left) and internal (right) smFRET pair labeled positions. Wild type (i.e., cysteine-containing) assays were performed as control separately for each labeling position. Error bars are from triplicate technical replicates.

**Fig S3.**
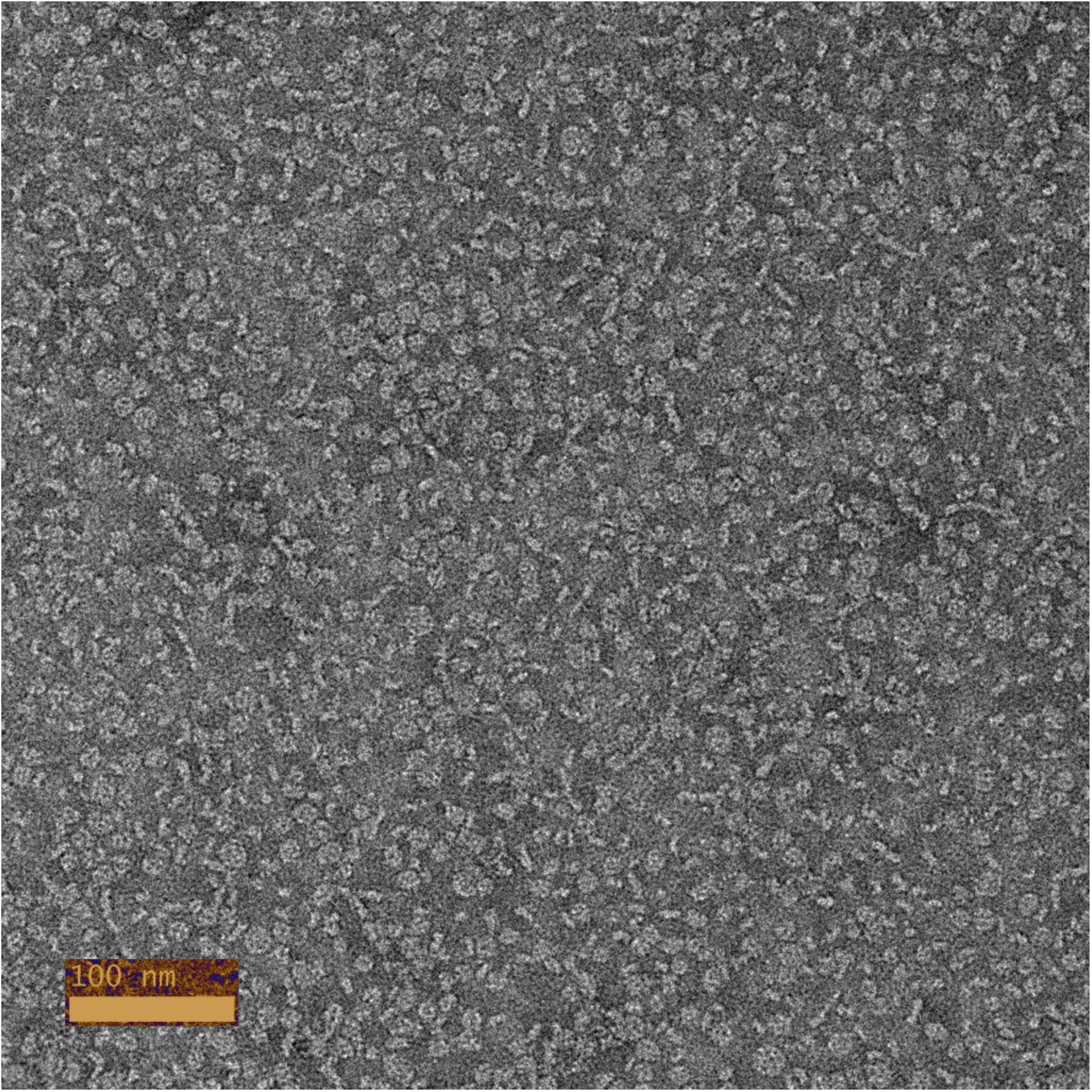
Negative stain electron microscopy of VcINDY reconstituted into lipid nanodiscs. Both top and side views of nanodiscs were seen. The nanodiscs had a diameter of ∼108 Å and each contained one VcINDY dimer. The sample was uniform and no aggregation was observed.

**Fig S4.**
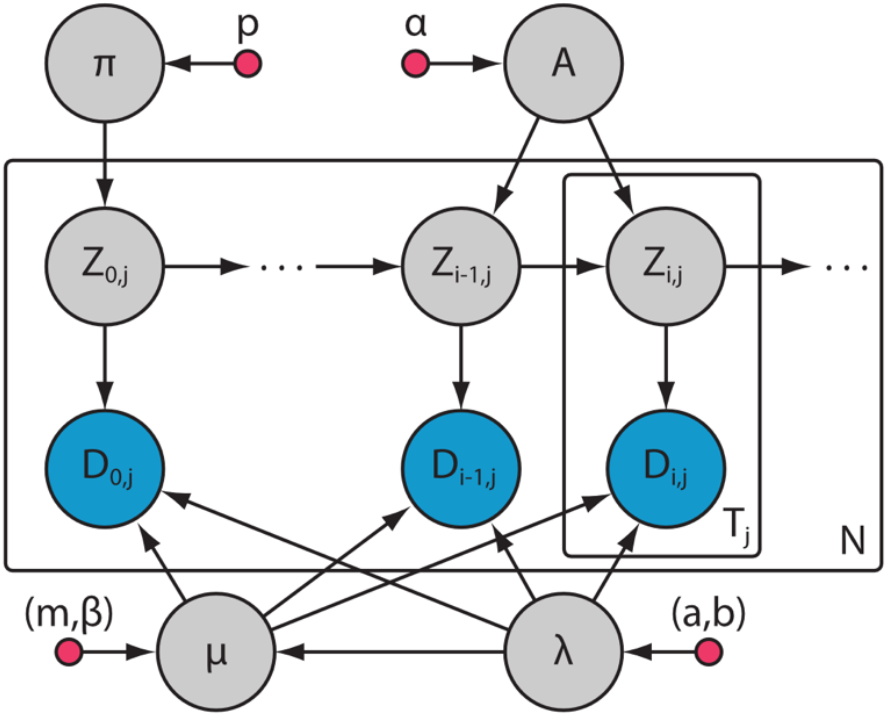
Directed acyclic graph for a global, variational Bayes HMM. Pink nodes represent hyperparameters, grey nodes represent unobserved variables, and blue nodes represent observed variables. T_j_is the number of data points in the j^th^ trajectory, and D_i,j_corresponds to the i^th^ data point of the j^th^ trajectory. Arrows represent conditional dependencies.

**Fig S5.**
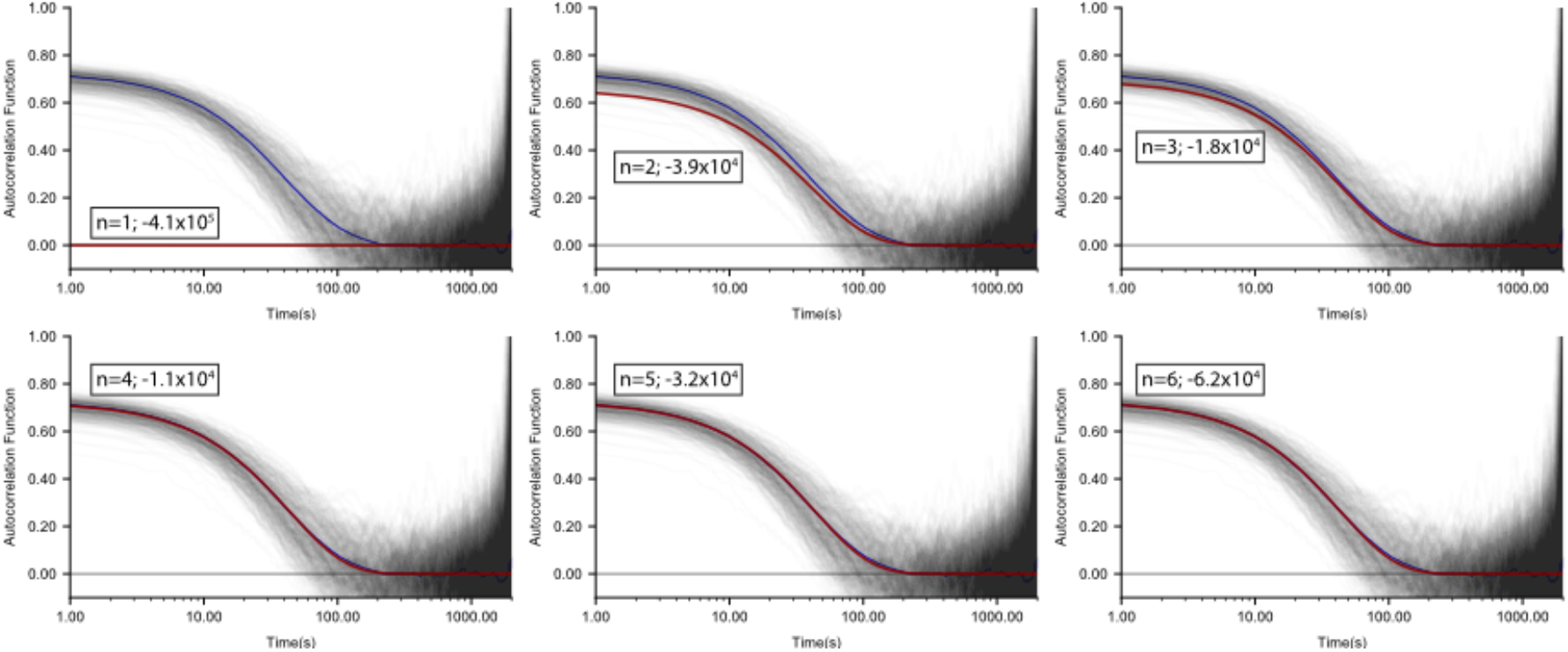
Autocorrelation function analysis of global HMMs with various number of hidden states of simulated, four-state E_FRET_ trajectories. Individual ACFs (black), ensemble ACF (blue), and global HMM ACF (red). The maximum ELBO HMM (n=4; ln(ELBO)= −1.1×10^4^) matches the number of E_FRET_ states in the simulation.

**Fig S6.**
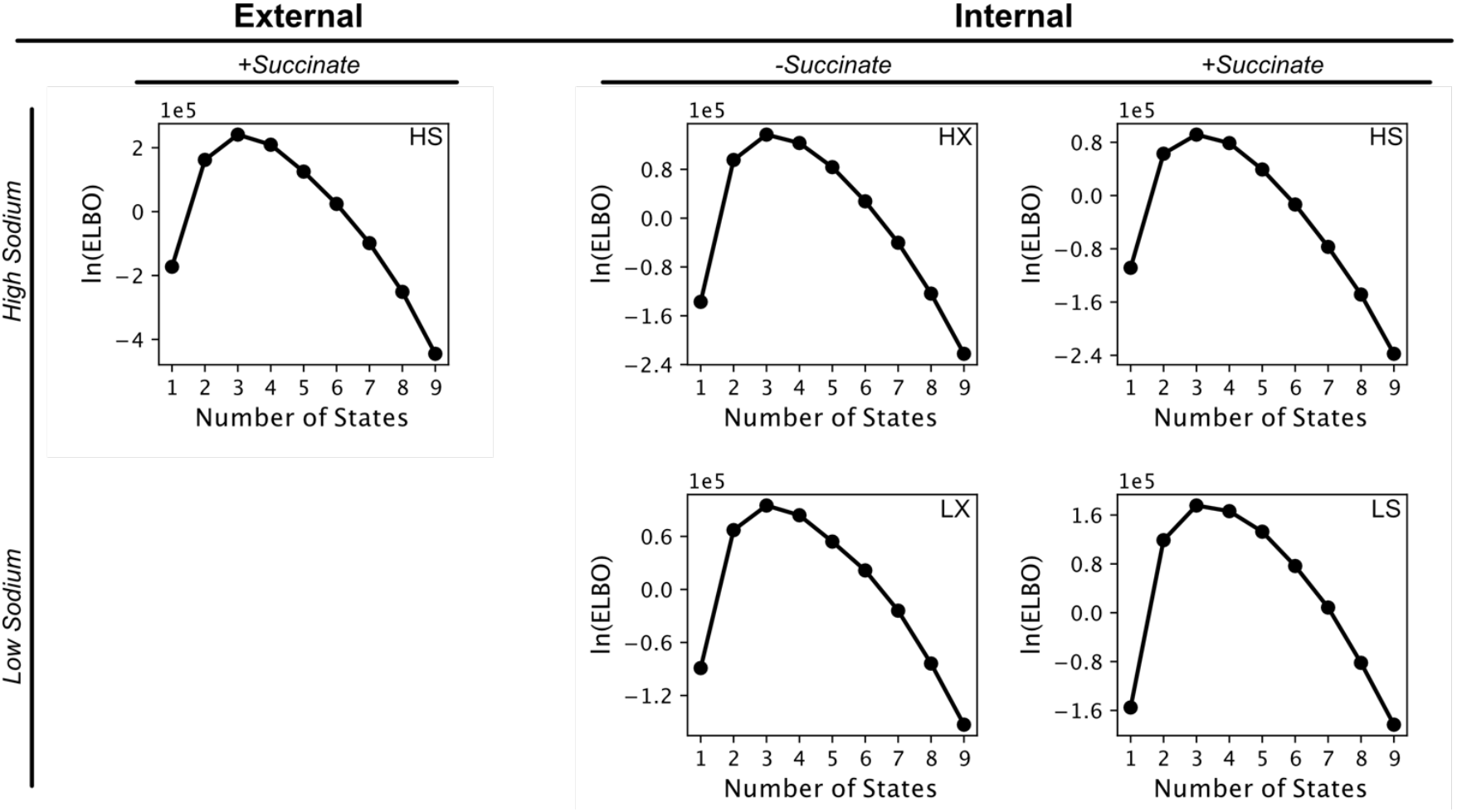
Substrate dependence of evidence lower bound (ELBO) calculations for global HMMs of VcINDY smFRET experiments. The maximum ELBO model is the HMM that best represents the set of E_FRET_ trajectories for those substrate conditions.

**Fig S7.**
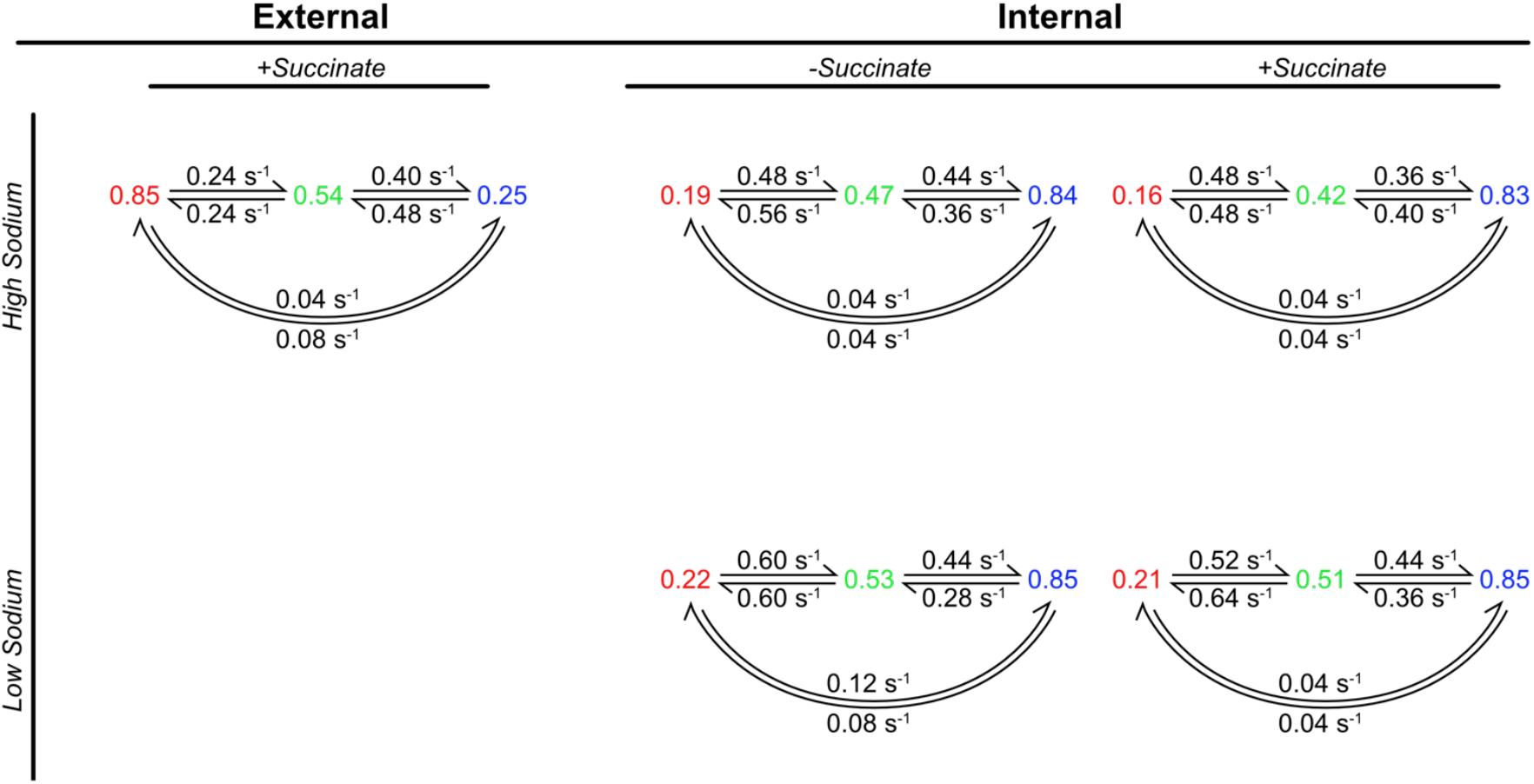
Substrate dependence of the HMM with maximum ELBO for global HMMs of VcINDY smFRET experiments. For each kinetic scheme, states are denoted by the mean E_FRET_ value of that state, which was inferred from the HMM. Rate constants for transitions between states are shown above the arrows.

**Fig S8.**
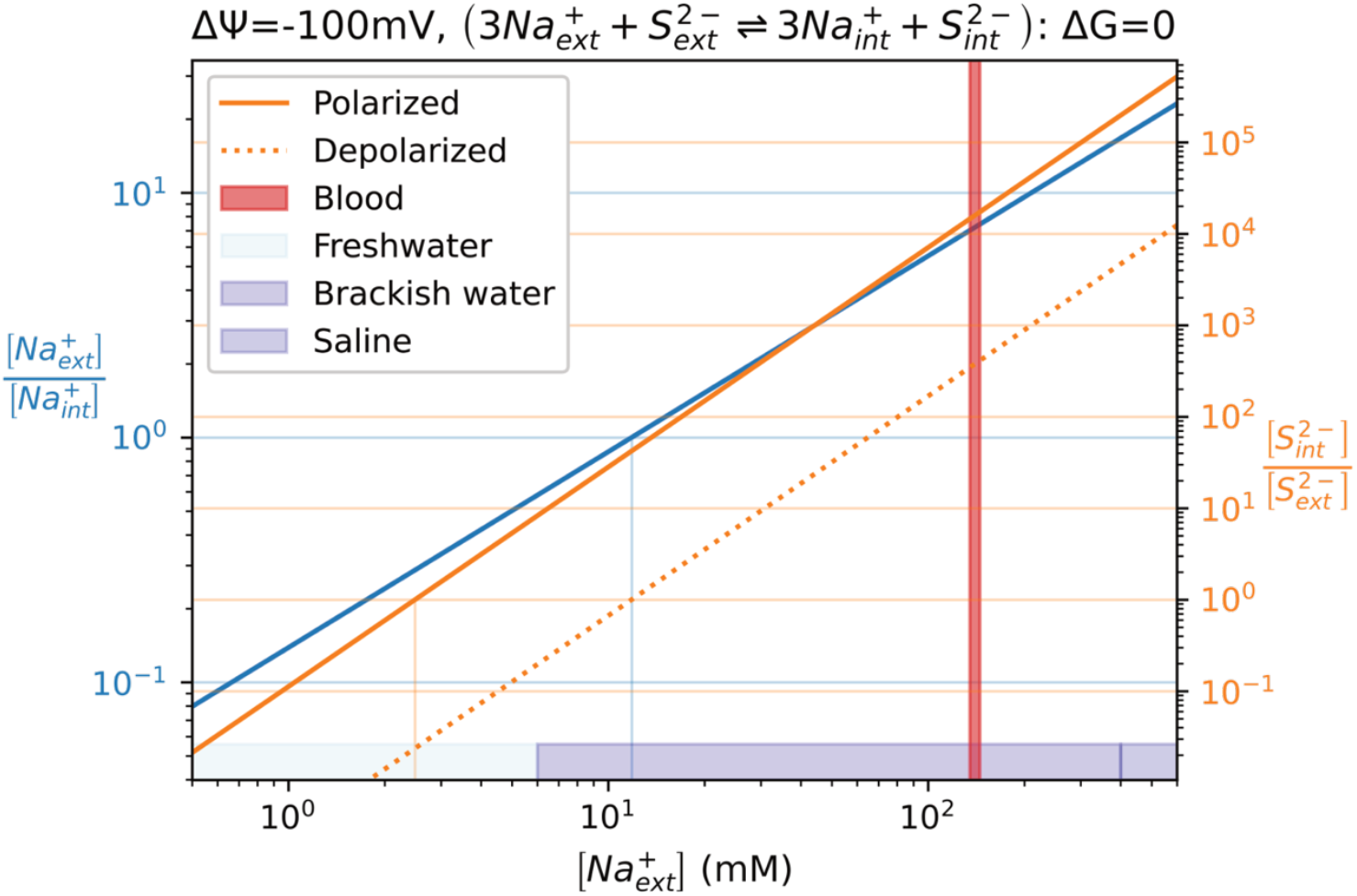
Estimated physiological Na^+^ and succinate concentrations in a bacterial cell in different environments. External (ext) and internal (int) Na^+^ concentrations were measured for another gammaproteobacteria, *E. coli*, at pH=7 (16). For *E. coli* at pH=7, the membrane potential difference (Δψ) is approximately -100 mV (17). With these numbers, balancing the chemical potential difference and effects of charge for the VcINDY transport reaction at equilibrium provides the internal to external ratio of succinate concentrations for the cell (orange). When Δψ = 0, the cell is depolarized (dashed).

**Fig S9.**
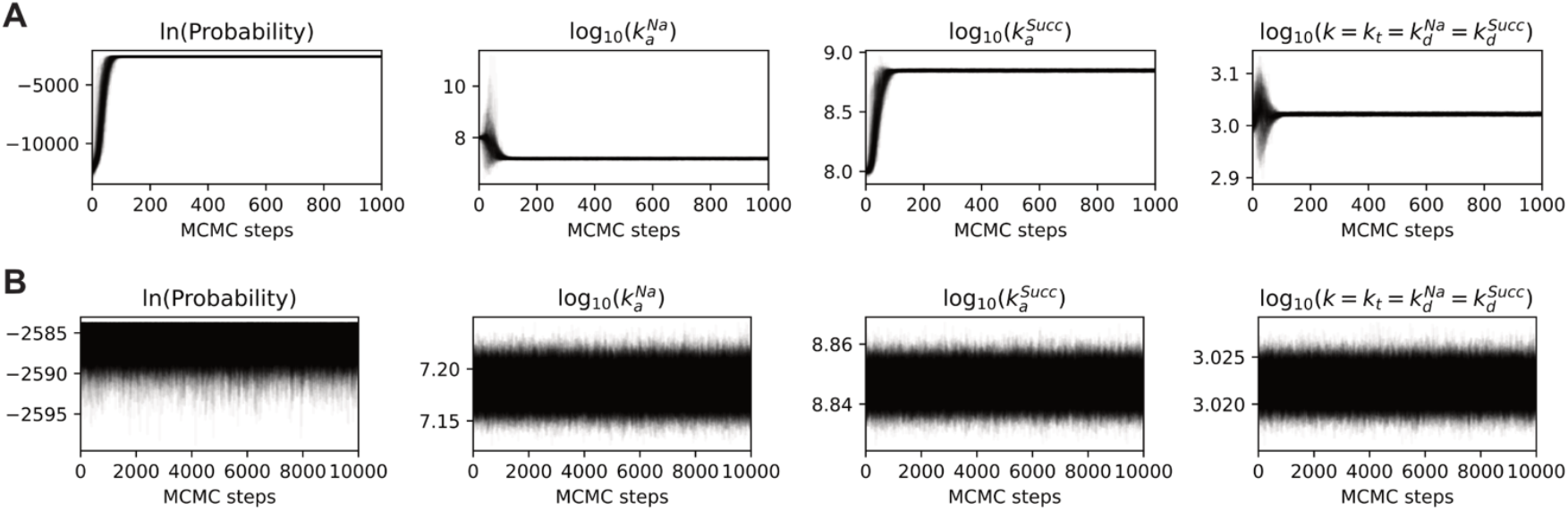
MCMC trajectories for optimal kinetic model parameters of VcINDY proteoliposome-based succinate transport assay. (A) Affine invariant ensemble MCMC sampler trajectories for 200 walkers during burn-in period. These samples were discarded. (B) Production period trajectories from 200 walkers. Only every 100^th^ sample of those shown here was used to minimize autocorrelation.

**Fig S10.**
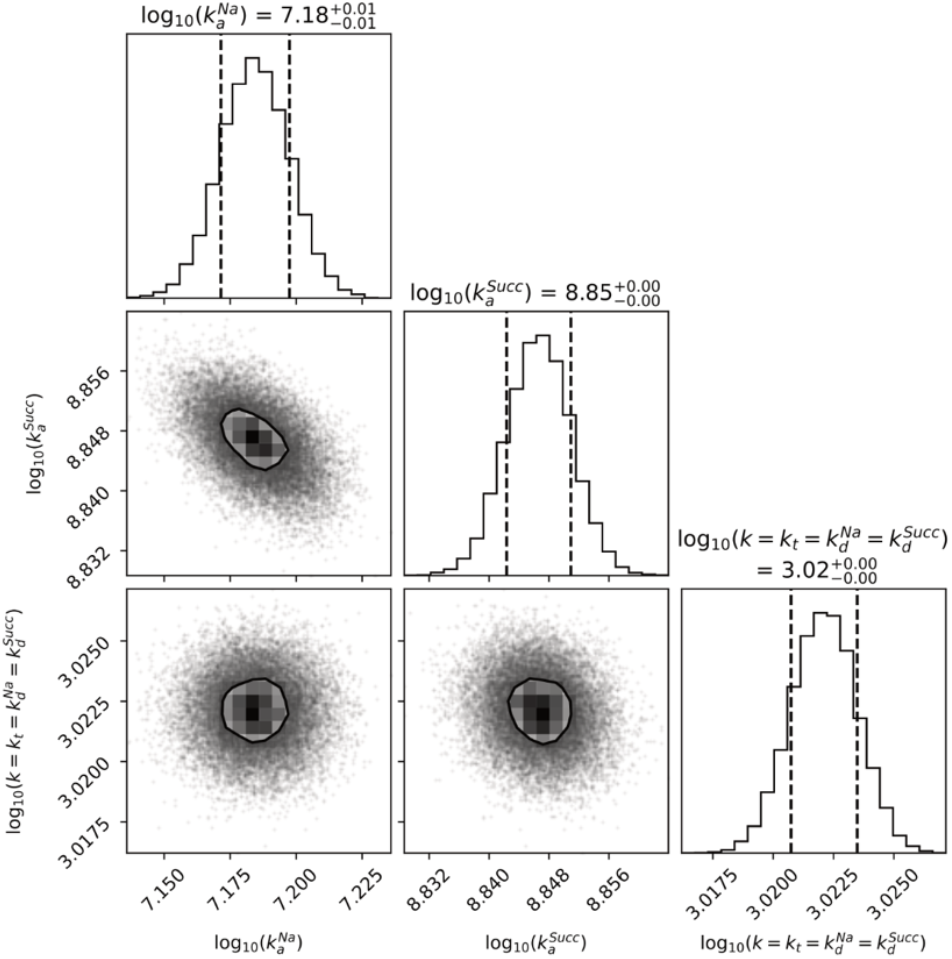
Corner plot of MCMC samples for optimal kinetic model parameters of VcINDY proteoliposome-based succinate transport assay. Ranges given are the 1σ level containing 39.3% of the sample volume.

**Fig S11.**
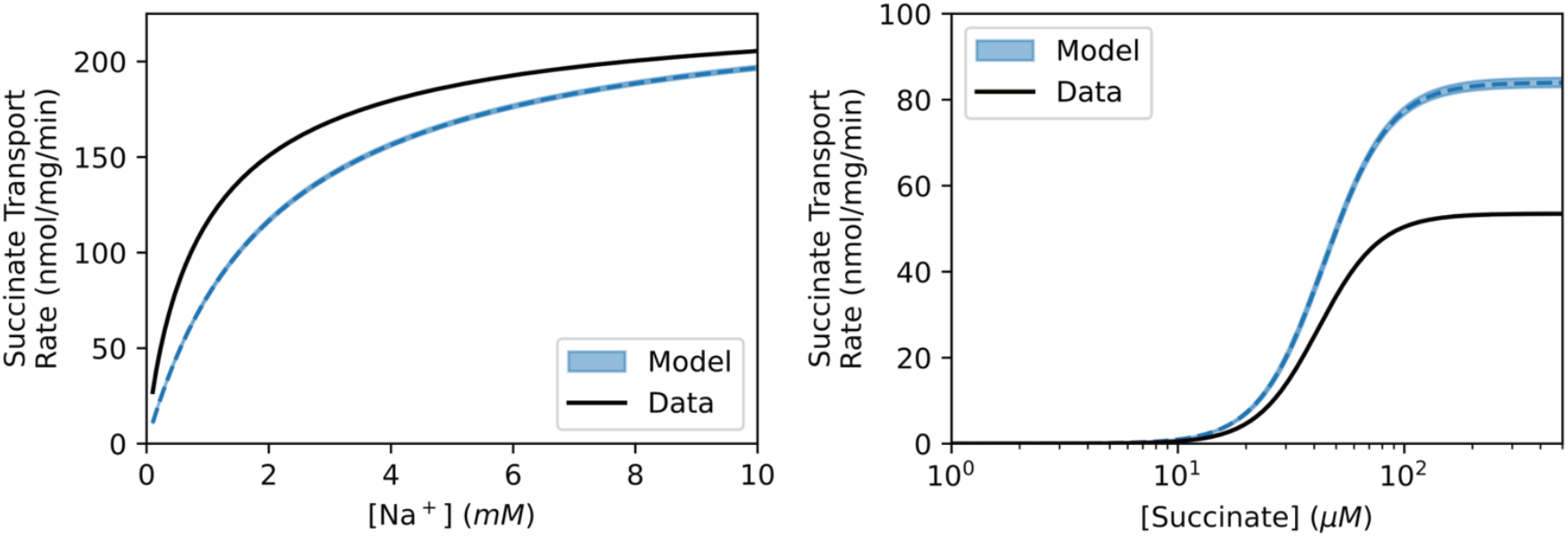
Modeled VcINDY transport rate experiments. Data from Mulligan and coworkers were reproduced from their reported *K*_*M*_, *V*_*max*_, and Hill coefficients (10). Modeled data are the 95% credible intervals generated from MCMC samples of 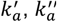.and ⟨*k*⟩, and dashed lines are the lowest energy solution.

**Fig S12.**
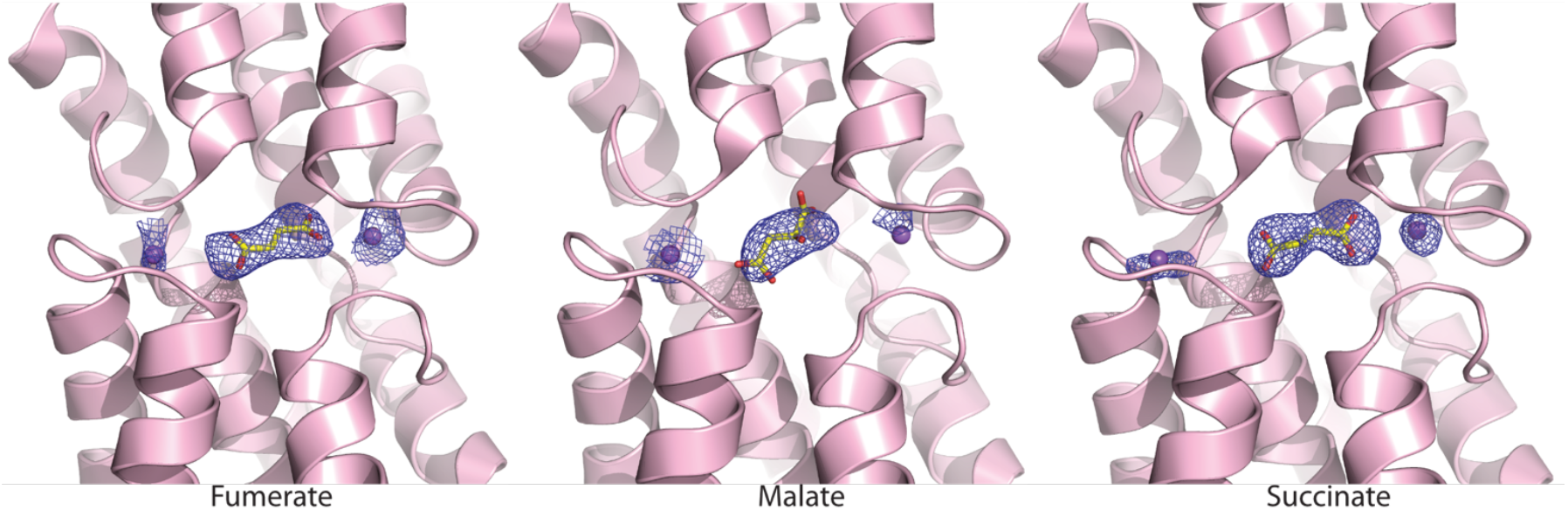
View of the VcINDY binding pocket from the cytosol. The X-ray crystallography derived structural model is of VcINDY (pink). Sodium ions are shown in purple, and specified dicarboxylate substrates are shown in yellow (carbon) and red (oxygen).

**Table S1.**
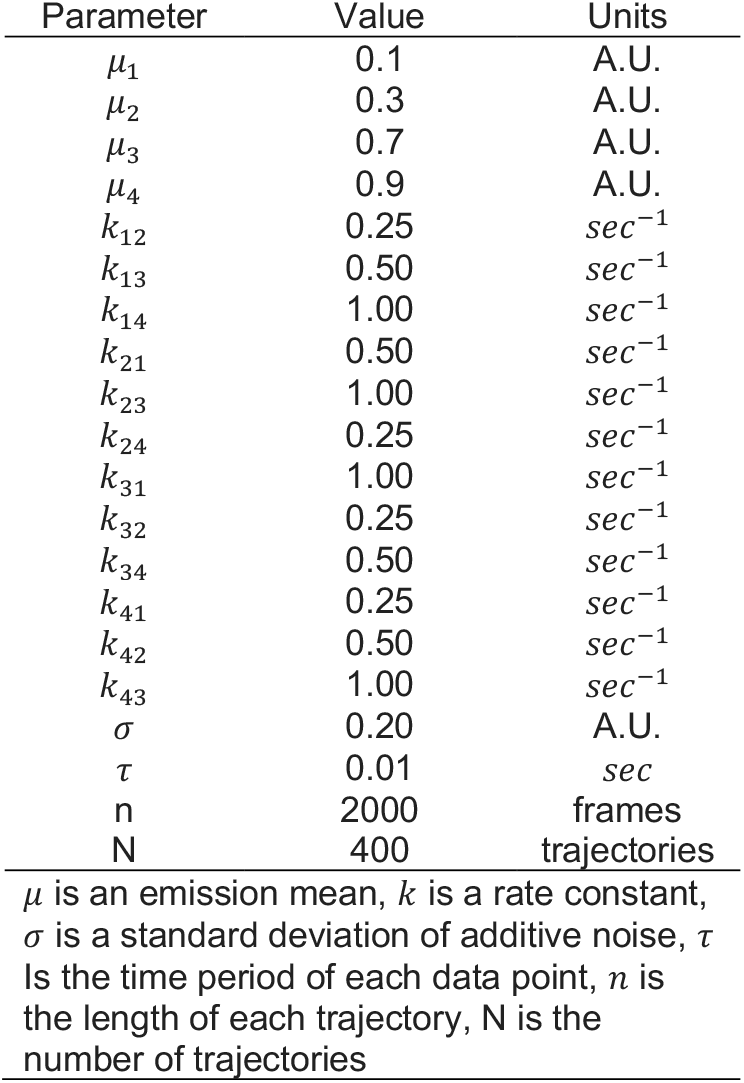
E_FRET_ trajectory simulation parameters.

**Table S2.** X-ray crystallography of VcINDY in complex with substrate.

| Parameter |  | VcINDY-succinate | VcINDY-fumarate | VcINDY-malate |
| --- | --- | --- | --- | --- |
| Data collection | Wavelength (Å) | 0.9791 | 0.9794 | 0.9794 |
|  | Resolution range (Å) | 50.0-3.09 (3.15-3.09) | 50-3.29 (3.35-3.29) | 50.0-3.50 (3.56-3.50) |
|  | Space group | P2<sub>1</sub> | P2<sub>1</sub> | P2<sub>1</sub> |
|  | Cell dimensions: a, b, c (Å) | 105.5, 102.5, 171.1 | 105.8, 102.1, 171.2 | 106.8, 102.4, 171.8 |
| | Cell angle: $\beta$ (°) | 98.808 | 98.937 | 97.791 |
|  | Total reflections | 128,621 (6,360) | 99,521 (3,598) | 86,871 (4,328) |
|  | Unique reflections | 65,778 (3,229) | 51,510 (1,913) | 44,866 (2,248) |
|  | Multiplicity | 5.0 (5.1) | 6.6 (5.3) | 6.9 (6.4) |
|  | Completeness (%) | 99.9 (100) | 93.9 (71.0) | 96.4 (96.9) |
| | $I/\sigma(I)$ | 10.3 (0.625) | 10.1 (0.667) | 10.8 (0.571) |
|  | Wilson B (Å<sup>2</sup>) | 80.97 | 110.64 | 134.20 |
|  | R<sub>pim</sub> | 8.4 (152) | 4.4 (54.2) | 3.5 (61.5) |
|  | CC<sub>1/2</sub> | 0.998 (0.443) | 0.102 (0.512) | 0.997 (0.604) |
| Refinement | R<sub>work</sub>/R<sub>free</sub> (%) | 25.3% / 29.6% | 24.9% / 28.5% | 26.1% / 30.5% |
|  | No. atoms - Protein/Ligands | 13,325 / 40 | 13,097 / 23 | 12,792 / 42 |
|  | RMSD bond lengths (Å) | 0.003 | 0.003 | 0.001 |
|  | RMSD bond angles (°) | 0.66 | 0.52 | 0.39 |
|  | Ramachandran favored (%) | 88.7 | 91.3 | 93.2 |
|  | Ramachandran allowed (%) | 9.99 | 7.30 | 5.72 |
|  | Ramachandran outliers (%) | 1.35 | 1.44 | 1.12 |
|  | Rotamer outliers (%) | 0.07 | 0.0 | 0.0 |
|  | Clashscore | 34.26 | 29.10 | 23.49 |
| Numbers in parentheses are statistics for the highest resolution shell. |  |  |  |  |

